# A quantitative genetics framework for understanding the selection response of microbial communities

**DOI:** 10.1101/2023.10.24.563725

**Authors:** Li Xie, Alex E Yuan, Wenying Shou

## Abstract

Heritability, a quantity that reflects the degree of resemblance between parent and offspring traits, is measured during plant and animal breeding because it predicts selection success during artificial selection of individuals. However, when whole microbial communities are under artificial selection to improve their traits, high heritability of the community trait does not necessarily predict selection success. To better understand the relationship between heritability and success during community selection, we establish a quantitative genetics framework, and in doing so, we obtain practical recommendations. Specifically, we decompose a community trait into “trait determinants”: genotype compositions and species compositions that impact the community trait and that vary among communities. This allows us to interpret heritability of a community trait in terms of the heritability of its determinants. We then use the Price equation to partition the selection response of a community trait into three phenomena: inter-community selection (heritability multiplied by selection intensity), transmission infidelity (the change in community trait from parent to offspring), and nonlinearity (due to a nonlinear relationship between parent and offspring traits). We illustrate that evolution within a community can cause the three terms to covary: in addition to the known effect of worsening transmission infidelity, intra-community evolution can lead to inflated heritability values greater than one (through an effect whereby “the poor get poorer”), and simultaneously magnify nonlinearity. As a consequence of these effects, heritability no longer predicts the selection response of a community trait. We propose effective selection strategies that improve heritability without accelerating intra-community evolution.

## Introduction

Microbes exist in communities of species that live together and interact. Microbial communities play fundamental roles in a variety of arenas, from human physiology to large-scale environmental systems. Engineered microbial communities could in principle make valuable contributions to medicine and industry. However, the functions performed by microbial communities often arise from a vast and largely uncharacterized web of interactions among member cells, making it difficult to apply rational design methods. Alternatively, artificial selection on whole communities represents a possible way to improve desirable traits without needing to decipher complex interactions [1].

In microbial community selection, we begin by initializing a population of “Newborn” communities by setting up each community with a starting amount of microbial cells. Individual cells in the communities then grow, interact, and possibly mutate for a period of time set by the experimentalist. After this time, the communities are said to have matured into “Adults” and are screened for the desired trait. The Adult communities with the highest trait values are then chosen for reproduction where each is used to inoculate multiple offspring Newborns for the next cycle. We will often refer to communities in one generation as “parents” and those of the next generation as “offspring”.

Selection of community traits has frequently proved challenging in both theoretical and experimental works. Selection responses can be unstable, unimpressive, or even opposite to the desired direction, and any improvements can quickly plateau [2, 3, 4, 5, 6, 7, 8, 9, 10, 11, 12]. A central question for researchers is: what factors determine the degree of success in a community-level artificial selection effort? Drawing from the vast literature on individual-level artificial selection, one idea is that selection campaigns will be more successful when a community trait has greater heritability [13, 14]. In a general sense, heritability refers to the extent to which related individuals resemble one another. There are several related (but nonequivalent) quantitative definitions for ‘heritability’ [15]. The biometric definition of heritability, the slope of the best-fit parent-offspring linear regression, is particularly informative here because it can be related to selection efficacy via a modified version of the Price equation.

Perhaps surprisingly, high heritability of microbial communities does not always predict effective selection [8], even when selection is applied to complex communities in which inter-community variation should arise readily [3]. To understand why, we need to consider the factors that influence the outcome of artificial selection, other than heritability.

The Price equation (Eq. 4 in Box 1) provides a convenient framework for studying the factors at play in artificial selection. This equation considers the intergenerational change in the average value of a trait (ΔAvg(*Z*)), and decomposes this quantity into three terms, each of which has a natural interpretation. The first term is trait heritability (*h*^2^) multiplied by the strength of selection for the trait, Cov 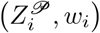. The importance of heritability and selection strength in artificial selection is self-evident. The second term (Avg (*δZ*_*i*_)), which we call transmission infidelity (or simply “infidelity”), represents systematic changes in trait values that may occur even in the absence of community-level selection. Infidelity might describe, for instance, the effect of a species whose high growth rate causes its frequency within each community to increase from one generation of communities to the next. The third term, Cov (*e*_*i*_, *w*_*i*_), describes the possible consequences of nonlinearity in the parent-offspring regression. Given that we have defined heritability as the slope of a line, it is sensible that if the true parent-offspring relationship is nonlinear, the slope alone may not be sufficient to describe intergenerational change. This term thus accounts for any nonlinear component of the parent-offspring relationship. We call this final term “nonlinearity”.

How important are infidelity and nonlinearity? Even in the context of individual-level selection, views in the literature differ widely. On one hand, as Helanterä and Uller [16] note, the infidelity and nonlinearity terms “are usually thought to be unimportant for adaptive evolutionary change, and dropped in most population genetic approaches”. If one does drop the infidelity and nonlinearity terms, the Price equation reduces to the breeder’s equation (Figure 1D; see also [17]). Conversely, some authors argue that such a simple linear relationship between heritability and selection efficacy is often an inaccurate description of biological reality [18, 19, 20]. Regardless, even if infidelity and nonlinearity are present, as long as they are uncorrelated with *h*^2^, we may still safely say that greater heritability is associated with more efficient selection.

**Figure 1:**
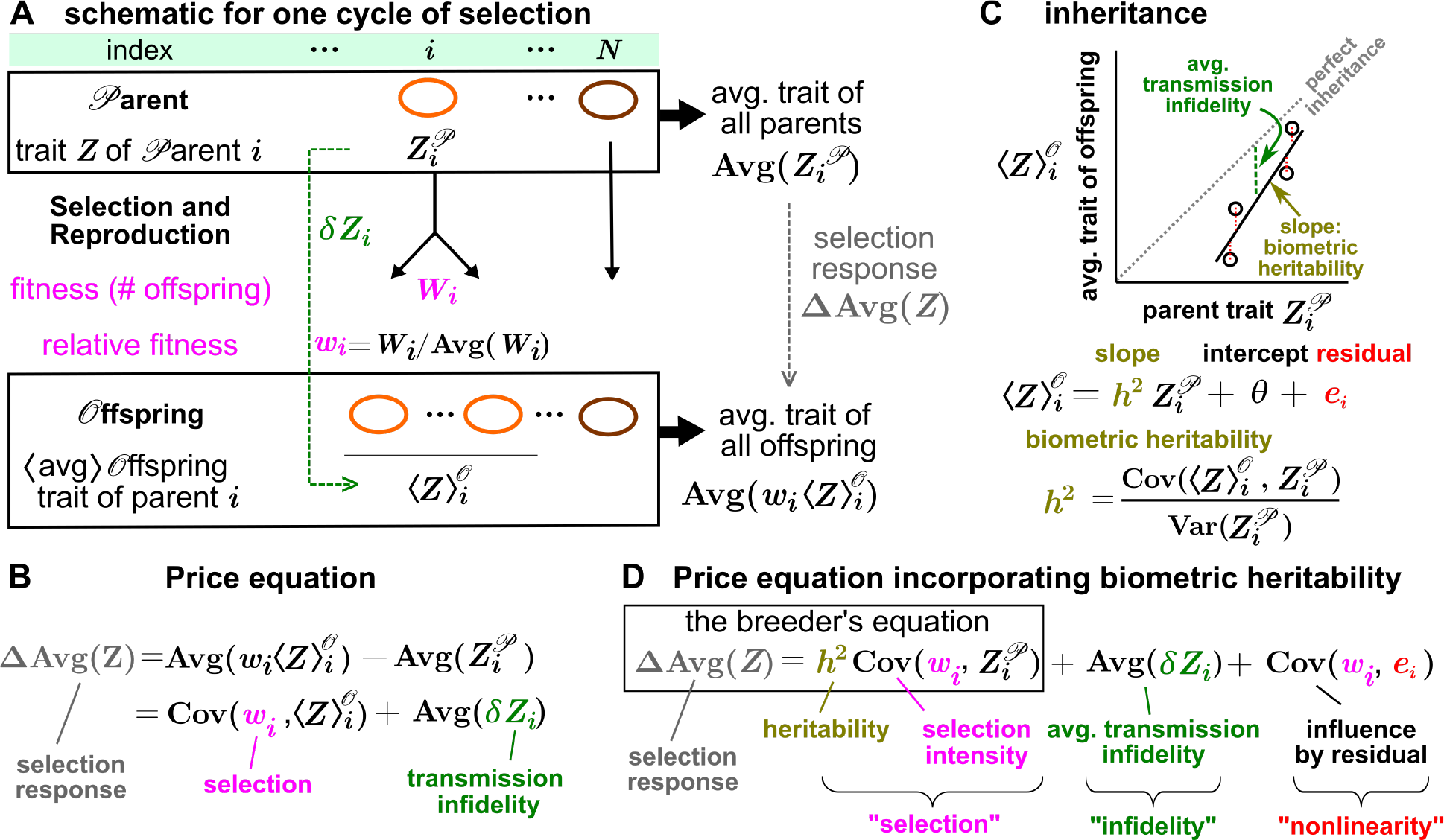
The Price equation is a general framework for describing selection response. (**A**) Schematic and nomenclature for one cycle of selection on *N* entities. **(B)** The Price equation describes selection response over one cycle. (**C**) Aspects of inheritance. Heritability (*h*^2^) is the slope of the parent-offspring regression. Infidelity (Avg(*δZ*_*i*_)) is the average change in trait from parent to offspring. Perfect inheritance (identical parent and offspring traits) occurs with aslope of 1 and infidelity of 0. (**D**) A modified Price equation that includes *h*^2^ and is an extension of the breeder’s equation.

In this article we show that the conventional wisdom of “higher heritablity begets better selection efficiency” does not hold in general for microbial communities. We begin with a motivating simulation example from our previously studied Helper-Manufacturer community [1, 21] where we observe correlations between heritability and selection response in some, but not other, settings. Next, to better understand the perhaps surprising decoupling between heritability and selection response, we develop a quantitative genetics framework for community-level selection. To do so, we introduce the concept of “determinants” for a community trait, and demonstrate that under certain conditions, a community trait within a selection cycle can be approximated as a linear function of trait determinants, similar to how an individual’s phenotype within a population can be approximated as a sum of contributions from genotype and environment. With this linear approximation, the heritability of a community trait can be understood in terms of the heritability of trait determinants. Using the trait determinant framework, we illustrate how intra-community evolution can increase the heritability of community traits, but negatively affect both the infidelity and nonlinearity terms of Eq. 4, thus offsetting the benefits from the increased heritability. We close by suggesting some selection strategies that may increase heritability without grossly accelerating intra-community evolution.

### Box 1

*A modified Price equation with an explicit heritability term*

Consider a population of *N* parent communities. Let the *i*th parent (*𝒫*) community have trait value 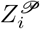 (Figure 1A). After selection on *Z*, the *i*th parent gives birth to *W*_*i*_ offspring, and thus has a fitness of *W*_*i*_ and relative fitness of *w*_*i*_ = *W*_*i*_*/*Avg(*W*_*i*_), where Avg(*…*) is an average across all parents. The *i*th parent’s transmission infidelity (given by 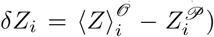 is the difference between average offspring trait 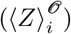 and parent trait 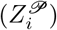. Here, *⟨…⟩* indicates an intra-lineage average, meaning 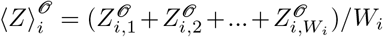 where 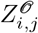 is the *j*th offspring of the *i*th parent. The Price equation (Figure 1B) gives the difference between the average offspring trait after selection 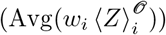 and average parent trait 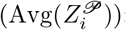:

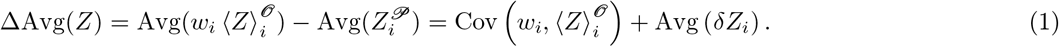

To write ΔAvg(*Z*) in terms of heritability, regress the offspring trait against the parent trait (Figure 1C):

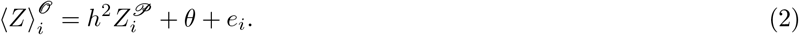

Here, *θ* is the intercept, *e*_*i*_ is the residual, and *h*^2^ (heritability) is the slope of the best-fit regression line:

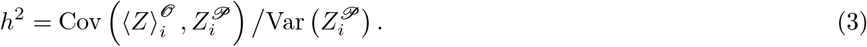

Substituting Eq. 2 into Eq. 1 then yields (Figure 1D):

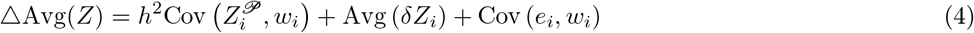

A more detailed derivation can be found in [14] or [22].

## Results

### Heritability predicts selection response in some contexts but not others

We examined the relationship between heritability and selection response using simulations of our previously studied [1, 21] Helper-Manufacturer (H-M) community model (Figure 2A) . In this community, H enables the growth of M by providing a Byproduct (at no cost to H) that is essential to M, and M manufactures a product (Product P) at a cost *f* to itself (where 0≤*f ≤*1). That is, M diverts a fraction *f* of its cellular resource to make Product instead of growing its own biomass. H and M also compete for a shared Resource. The community trait of interest, denoted *P* (*T*), is the total amount of Product accumulated by the end of the maturation time *T* . The community trait requires both H and M, and is costly to M. Note that costly community traits such as *P* (*T*) are common: engineered microbes must divert cellular resources to make a drug designed by the experimentalist; if multiple species are required for a community trait, fast-growing species must evolve slower growth to avoid outcompeting their partners.

**Figure 2:**
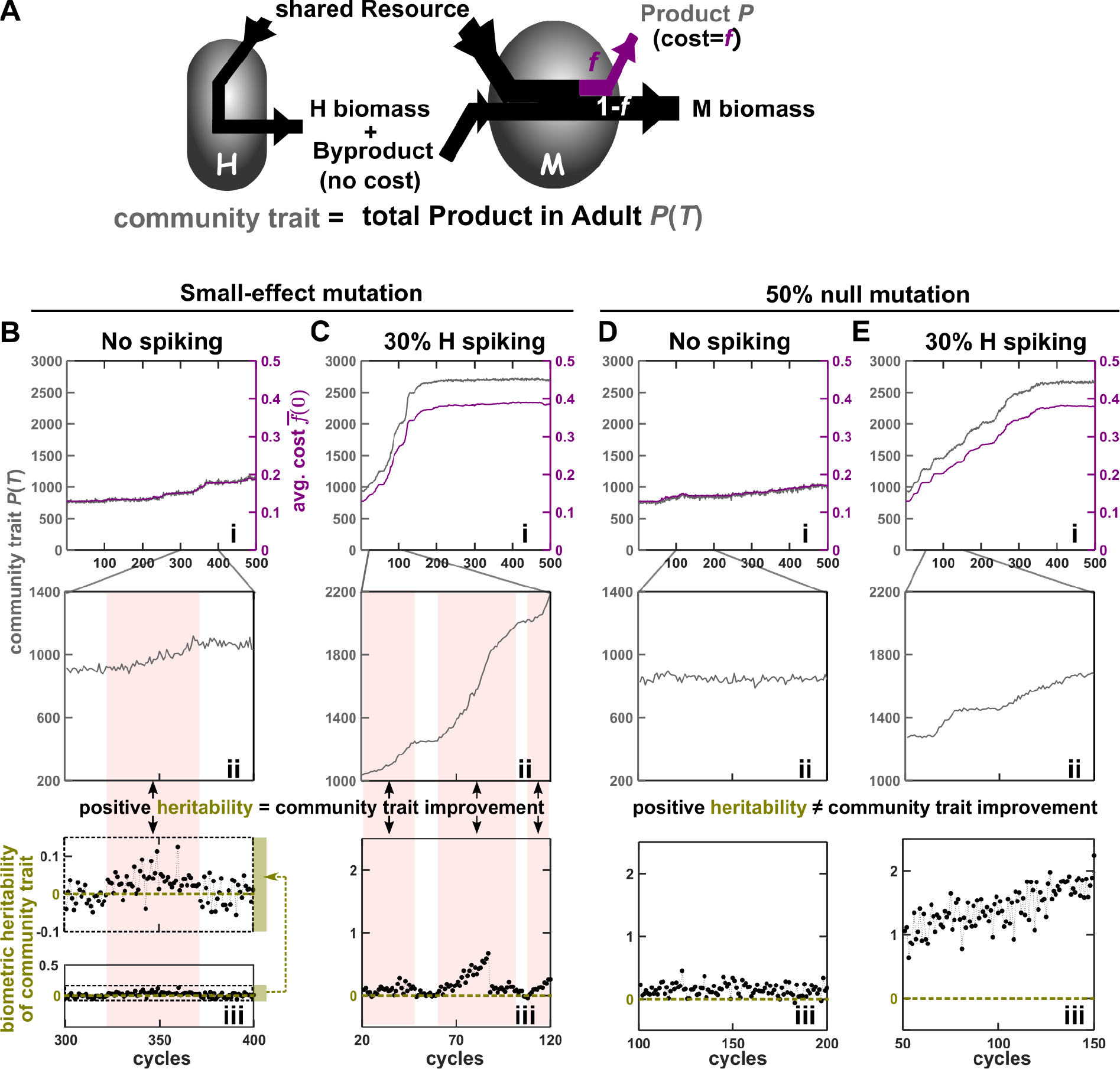
Biometric heritability predicts selection response in some contexts but not others. (A) Schematic of the H-M community. H-M community selection was carried out under different reproduction strategies (no spiking in (B, D) vs 30% H spiking in (C, E)) and mutation spectra (small-effect mutations in (B, C) vs 50% null mutations in (D, E)). B(i)-E(i): evolutionary dynamics of over 500 cycles. The community trait, *P* (*T*), is shown in grey. The biomass-weighted intra-community average of the Newborn cost parameter, 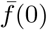, is shown in purple. Both the *P* (*T*) and 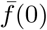 curves show an average of the respective quantity among selected communities only. The dynamics over 100 cycles are expanded in B(ii)-E(ii). The biometric heritability over the same 100 cycles are plotted in B(iii)-E(iii). To quantify biometric heritability of community trait, each parent Adult reproduces 10 offspring. When the average offspring community trait is regressed against the parent community trait, the slope is the biometric heritability. In Band C, pink blocks mark regions where heritability is on average positive, and only in these regions community function improves. Higher heritability of community trait leads to faster improvement within the same mutation spectrum (C faster than B ; E faster than D), but not necessarily across different spectra (e.g. compare D with B; E with C). All graphs are plotted on the same scale, except the top panel of B(iii) which is a magnified view of the bottom panel.

In our individual-based stochastic simulations of community selection, each cycle starts with 100 Newborn communities. Each Newborn is initiated with the same total species abundance (total biomass fixed to 100, which is equivalent to 50∼100 cells each with a biomass between 2 and 1) and the same amount of Resource (supporting 10^4^ cells or 6∼7 doublings). The low population size and low number of doublings collectively ensure that evolution is slow within a cycle, which will be helpful later in satisfying a key assumption of our mathematical approach (Assumption 1 in Box 2). Communities are reproduced by simulated pipetting so that the species proportions in a Newborn fluctuates around that of the parent Adult. Among genotype values, only M’s cost *f* is allowed to mutate. When an M cell divides, each daughter has a probability of to change its cost *f* . We compare two mutation spectra: “small-effect mutations” (randomly increasing or decreasing cost *f* by on average 0.01) and “50% null mutations” (50% of mutations reducing *f* to 0, and the rest being small-effect). Intra-community evolution is faster in the presence of null mutations because they confer the maximal growth advantage. At the end of maturation, we choose the 10 Adult communities exhibiting the highest total Product, and allow each Adult to reproduce 10 Newborn communities [1]. See Supplement 6 for more details.

Besides comparing two mutation spectra (“small-effect” versus “50%-null”), we also compare two reproduction strategies [21]: In the “no-spiking” strategy, H and M cells in a parent Adult are randomly assigned to each Newborn until a total of 100 biomass units is reached. In the “30%-H spiking” strategy, an offspring Newborn obtains 70% of its biomass from its parent Adult and the other 30% from a reservoir of H cells.Note that since the only mutable parameter is M’s *f*, the H cells do not evolve. In both strategies, the fraction of M biomass in offspring Newborn fluctuates stochastically to reflect random sampling of the parent Adult (with a coefficient of variation around 10%∼30%). As shown previously [21], the heritability of the community trait is higher when Adult communities are reproduced with 30%-H spiking than with no spiking strategy (Figure 2iii, C higher than B; E higher than D).

Within the same mutation spectrum, higher heritability of the community trait is associated with greater selection response. Specifically, the 30%-H spiking strategy leads to a faster increase in the community trait *P* (*T*) and in the average cost parameter 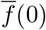 (Figure 2i, C faster than B; E faster than D). Similarly, the 30%-H spiking strategy leads to a faster increase in the average cost parameter at *T* = 0, as shown in purple in Figure 2B(i)-E(i).

However, when we compare segments of simulations under different mutation spectra, greater heritability no longer indicates greater selection response. Specifically, the segments from the 50%-null mutation spectrum have greater heritability than their small-effect spectrum counterparts (Figure 2D(iii) higher than B(iii); E(iii) higher than C(iii)), but lower selection response (Figure 2B(ii) vs D(ii); C(ii) vs E(ii)).

### Changes in the selection term can be coupled to opposing changes in the infidelity and nonlinearity terms

To further explore the relationship between heritability and selection response, we revisited the time segments in Fig 2 through the lens of the Price equation. For each segment, we estimated each of the three terms in the Price equation (selection, infidelity, and nonlinearity) by simulating the offspring trait values for all communities at each cycle, including non-selected communities. Consistent with Fig 2B(iii)-E(iii), the selection term (which contains heritability) is indeed larger under the 50%-null mutation spectrum (Fig 3, compare A vs C; B vs D). However, the time segments from the 50%-null mutation spectrum also have a more strongly negative transmission infidelity term (Fig 3; same panel comparisons). In the 30%-H spiking condition, the time segment in with the 50%-null mutation spectrum additionally has a more strongly negative nonlinearity term (Fig 3, compare B vs D).

**Figure 3:**
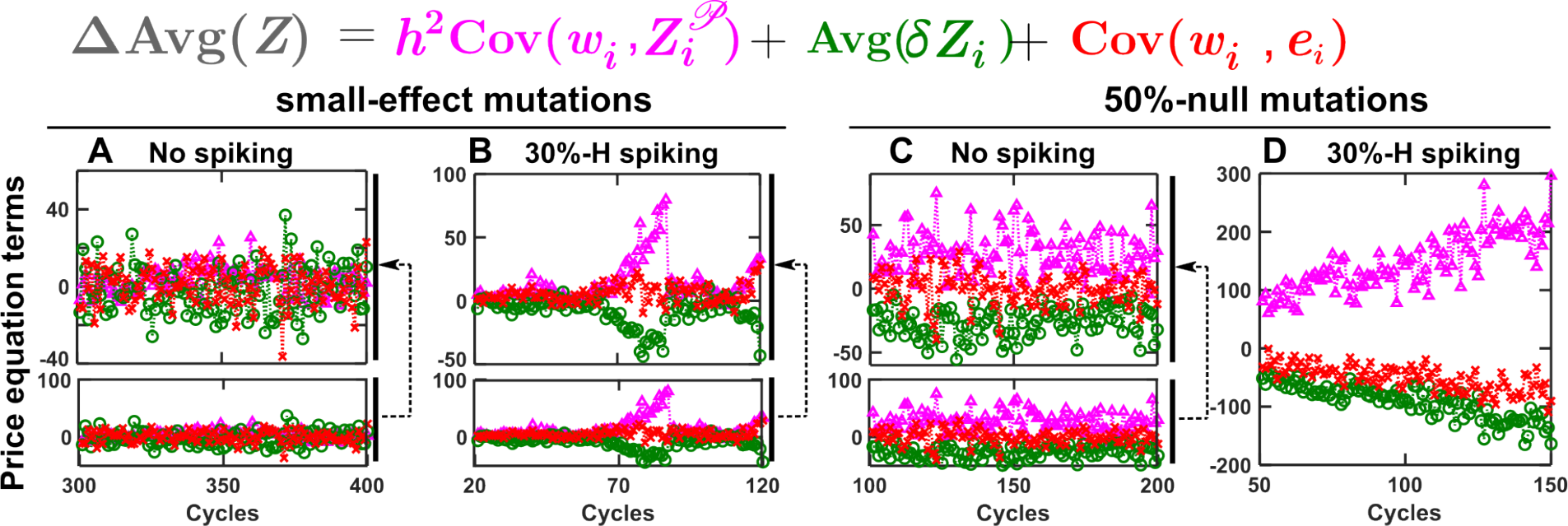
The three terms in the Price equation can exhibit correlated dynamics. The three terms of the Price equation (selection term in magenta triangles, infidelity term in green circles, and nonlinearity term in red crosses) over 100 cycles in Figure 2B(iii)-E(iii) are plotted in A-D, respectively. The presence of null mutations in the population leads to simultaneously a larger selection term and a more strongly negative infidelity term (compare C vs A, and D vs B). Additionally, in the condition with null mutations and 30%-H spiking (D), the increase in the selection term coincides with a decrease both the infidelity and nonlinearity terms.In Figure S3 we directly plot the selection term against the infidelity and nonlinearity terms, and test for the statistical significance of the correlations in an autocorrelation-aware way. All graphs are plotted on the same scale, except for the top panels of A-C which are magnified from the bottom panels.

We also observed that changes in the selection term could be coupled to countervailing changes in infidelity and nonlinearity across cycles (Fig S3). Specifically, we observed a negative correlation between the selection term and infidelity in all cases except the condition with no spiking and small-effect mutations (Fig S3B-D). We also observed a negative correlation between the selection term and nonlinearity under the 50%-null mutation spectrum, regardless of the spiking condition (Fig S3C-D).

The results so far indicate that under some conditions, an increase in the selection term of the Price equation can be coupled to opposing changes in the infidelity and/or nonlinearity terms, thereby cancelling or even overturning the overall change in selection efficacy. When should we expect a coupling between the selection term and the other terms of the Price equation? To address this question, it will he be helpful to draw on an approach from quantitative genetics that views a phenotypic trait as the sum of environmental and genetic factors. In the following, although we seek general lessons, we will refer to the H-M community as a specific case study. Additionally, although the term “community” typically refers to assemblages of two or more species, our theoretical development will include group-level selection in monocultures as a special case.

### A community trait can be approximated as a linear function of trait determinants under certain assumptions

Traits of a community emerge from interactions between individuals, similar to traits of an individual that emerge from interactions between the genotype and environment. In classic quantitative genetics of individual traits, linear models form the backbone of analysis [23, 24]. A trait is often modeled as a sum of contributions from genotype and environment (Eq. S8). The genotype effect may be further decomposed into a sum of terms such as additive effects of alleles, dominance effects (i.e. interactions between alleles at a givenlocus), and epistatic effects (i.e. interactions among loci). For example, the sheep litter size can be modeled as a linear function of the specific alleles of the Booroola gene and any dominance effects [23]. As another example, the nitrate reduction rate of a proteobacterium can be modeled as a linear function of the presence or absence (1 or 0) of relevant genes [25].

Similarly, community traits can be understood in terms of *community trait determinants* (“determinants”): factors that vary among communities and whose variation can cause variation in the community trait [21]. To limit the number of determinants, we assume that the initial abiotic factors are identical across all Newborn communities. If a community trait is not heavily influenced by stochastic events that occur during community maturation (e.g. newly arising mutations), then the community trait is primarily determined by species abundances and phenotypes in the Newborn. The initial species abundances in a Newborn community are defined as *species abundance determinant* s, since these can vary among Newborns when, for example, small samples are taken from an Adult community to seed Newborns via a pipet. Within a species, for each phenotype that varies among communities and that affects the community trait (e.g. growth rate, affinity for metabolites), a *genotype value* can be assigned to an individual, and the *average* genotype value in a Newborn is defined as a *genotype determinant* of the community trait. The average genotype values can adequately predict a community trait as long as the intra-community genotype value distributions are tight and remain nearly constant within one maturation cycle (Assumption 1 in Box 2). For example, consider an exponentially growing population consisting of *M* (0) individuals with different growth rates (different shading in Fig 4A), and no mutations. The community trait of interest — the population size at time *T* (*M* (*T*)) — is the sum of the population sizes of all *M* (0) lineages (Fig 4A). When Assumption 1 is satisfied, the community trait (*M* (*T*)) can be predicted by multiplying the species abundance determinant (Newborn population size *M* (0)) with an exponential function of the genotype determinant (average growth rate at the Newborn stage 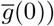 (Figure 4B; Figure 4D(i)). If Assumption 1 is violated, the prediction is poor (Figure 4D(ii)).

**Figure 4:**
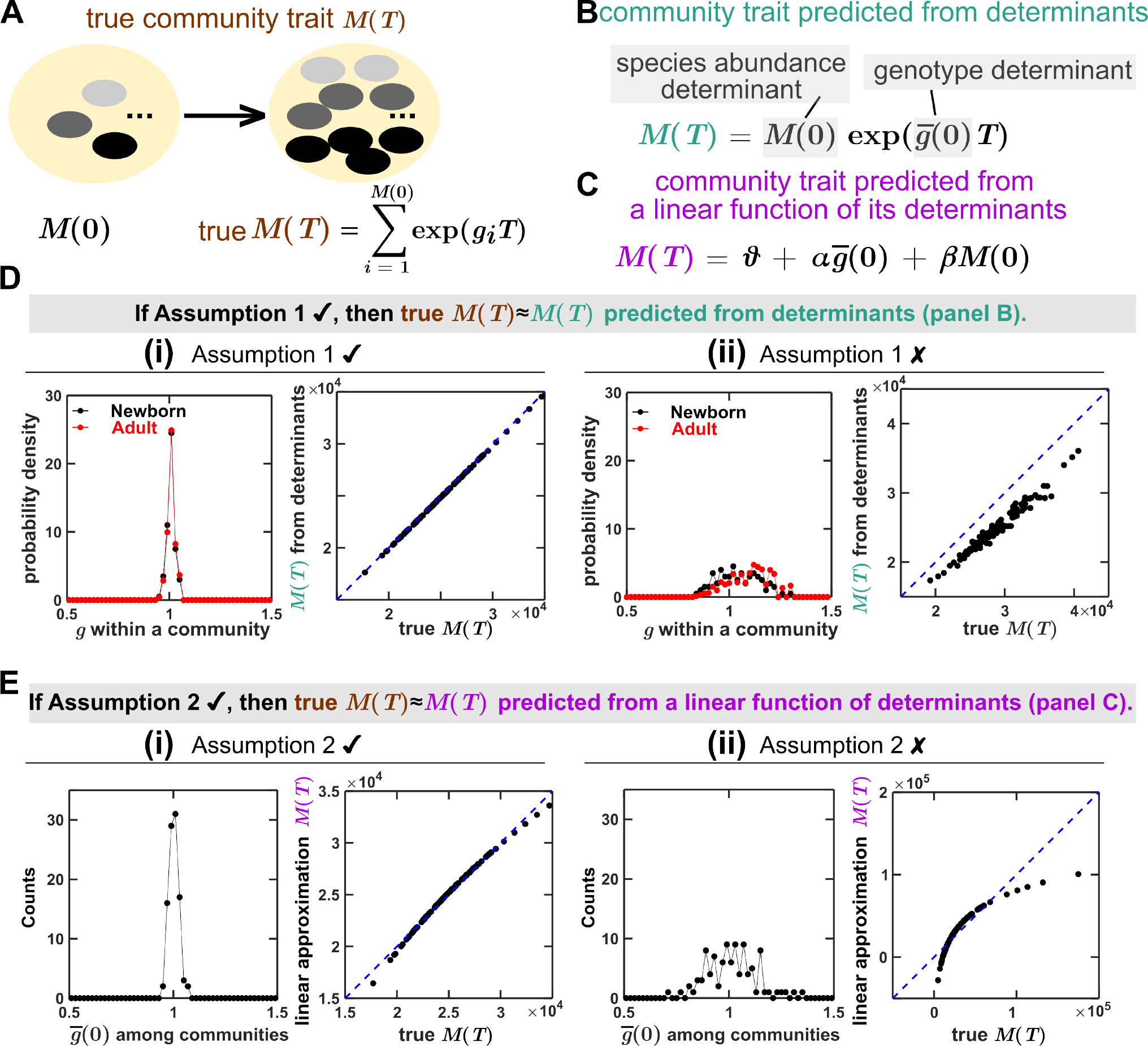
An example where a community trait can be approximated as a linear function of determinants. Here, we consider *N* = 100 communities in unrestricted nutrient. For simplicity, each Newborn community has *M* (0) = 100 microbes, and thus species abundance is not a determinant (as it is fixed across communities). The community trait is *M* (*T*), the total population size of an Adult community at *T* = 5.5 (∼8 doublings). (**A**) When a community (yellow oval) is not clonal, 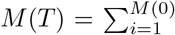 exp(*g*_*i*_*T*) is the true community trait, where *g*_*i*_ (indicated by the grayscale of small oval) is the growth rate of the *i*th microbe in the Newborn. **(B, D)** If the growth rates of individuals within a Newborn community are tightly distributed, then the true community trait can be accurately predicted from the average growth rate determinant 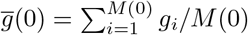, i.e., 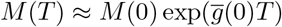. The genotype value *g* of 100 individuals in a Newborn community are drawn from a normal distribution with standard deviation of 0.02 in D(i) and 0.1 in D(ii). Note that when the distribution of *g* is narrow, it does not change from Newborn (black) to Adult (red). But when the distribution of *g* is wide, it shifts to the right over time as fast growers increase in frequency. (**C, E**) Furthermore, if the inter-community distribution of the determinant 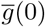 is narrow, then community trait *M* (*T*) can be approximated as a linear function (the blue dashed line) of the genotype determinant 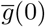. The genotype determinant 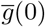 of 100 Newborn communities are drawn from a normal distribution with standard deviation of 0.02 in E(i) and 0.1 in E(ii).

If we additionally assume tight distributions of determinants among communities within one cycle (Assumption 2 in Box 2), a community trait can be further approximated as a *linear* function of determinants. This is based on the intuition that any differentiable curve, when closely observed over a narrow range, resembles a straight line. Going back to our previous example, if Assumption 2 is satisfied (e.g. the growth rate determinant 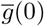 is tightly distributed among all communities), then we can approximate community trait *M* (*T*) as a linear function of the two determinants (Figure 4C; Figure 4E(i)). Otherwise, the linear approximation is poor (Figure 4E(ii)). The mathematical details for this example, and a more complex example of three-species multi-genotype communities, can be found in Supplement 1.

#### Box 2

*Approximating a community trait as a linear function of determinants: two assumptions*

1. *Sufficiently slow intra-community evolution: Within a community, genotype values of individuals within a species exhibit a narrow distribution* ***and*** *remain nearly constant from Newborn to Adult*. This assumption allows us to use the average genotype values (rather than the entire distribution) in a Newborn community (rather than throughout maturation) as determinants to predict a community trait. To satisfy this assumption, evolution must be sufficiently slow. This in turn requires the mutation rate, the fitness differences between genotypes, the population size, and the number of generations during each cycle of community maturation to be sufficiently small.
2. *Among communities from one selection cycle, distributions of determinants are sufficiently narrow*. This is valid when inter-community selection is sufficiently strong. Even though a community trait is generally a nonlinear function of determinants, if the ranges of determinants are sufficiently small, the linear approximation is valid.

### The Helper-Manufacturer community trait can be approximated as a linear function of a genotype determinant and a species abundance determinant

Let us return to the Helper-Manufacturer (H-M) community (Figure 5A). The community trait *P* (*T*) has two determinants, a species abundance determinant and a genotype determinant. Although the H-M community has two species, since the total Newborn biomass is fixed, only one determinant is needed to describe species composition: the proportion of a Newborn’s biomass that is due to M cells, denoted *ϕ*_*M*_ (0). Since all genotypes (except for M’s cost *f*) are constant by construction, the only genotype determinant is the average cost of M in Newborn, denoted 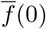. Although the community trait *P* (*T*) is a nonlinear function of the two determinants (Figure S1), we find that within a selection cycle, it can be approximated as a linear function of its two determinants (Figure 5A):

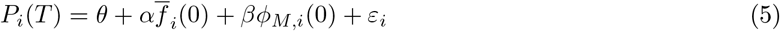

Here, the index *i* denotes the *i*th community, the coefficients *α* and *β* indicate the contributions of the two determinants to the community trait, *θ* is the intercept, and *ϵ*_*i*_ is the residual of a least-squares linear regression. To simplify notation, we define for the *i*th community its genotype determinant 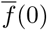 as *x*_*i*_, its species abundance determinant *ϕ*_*M*_ (0) as *y*_*i*_, and its community trait *P* (*T*) as *Z*_*i*_. Eq. 5 may be rewritten:

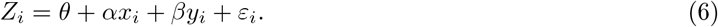

In general, if the two assumptions (Box 2) are valid, then the trait of *i*th community can be reasonably approximated by a linear combination of its determinants:

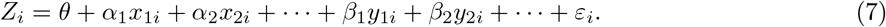

where *x*_1*i*_, *x*_2*i*_,…, are genotype determinants of genotypes 1, 2… of various species in the *i*th Newborn community; *y*_1*i*_, *y*_2*i*_, … are species abundance determinants in the *i*th Newborn community, *θ* is the intercept, and *ϵ*_*i*_ is the residual. Although the *α* and *β* coefficients can change over the long term, if Assumption 1 (slow evolution) is satisfied, they will remain nearly constant between one cycle and the next. In Supplement Section 1, we show another example where the total biomass trait of a three-species competitive community has a total of 18 species abundance determinants and genotype determinants.

**Figure 5:**
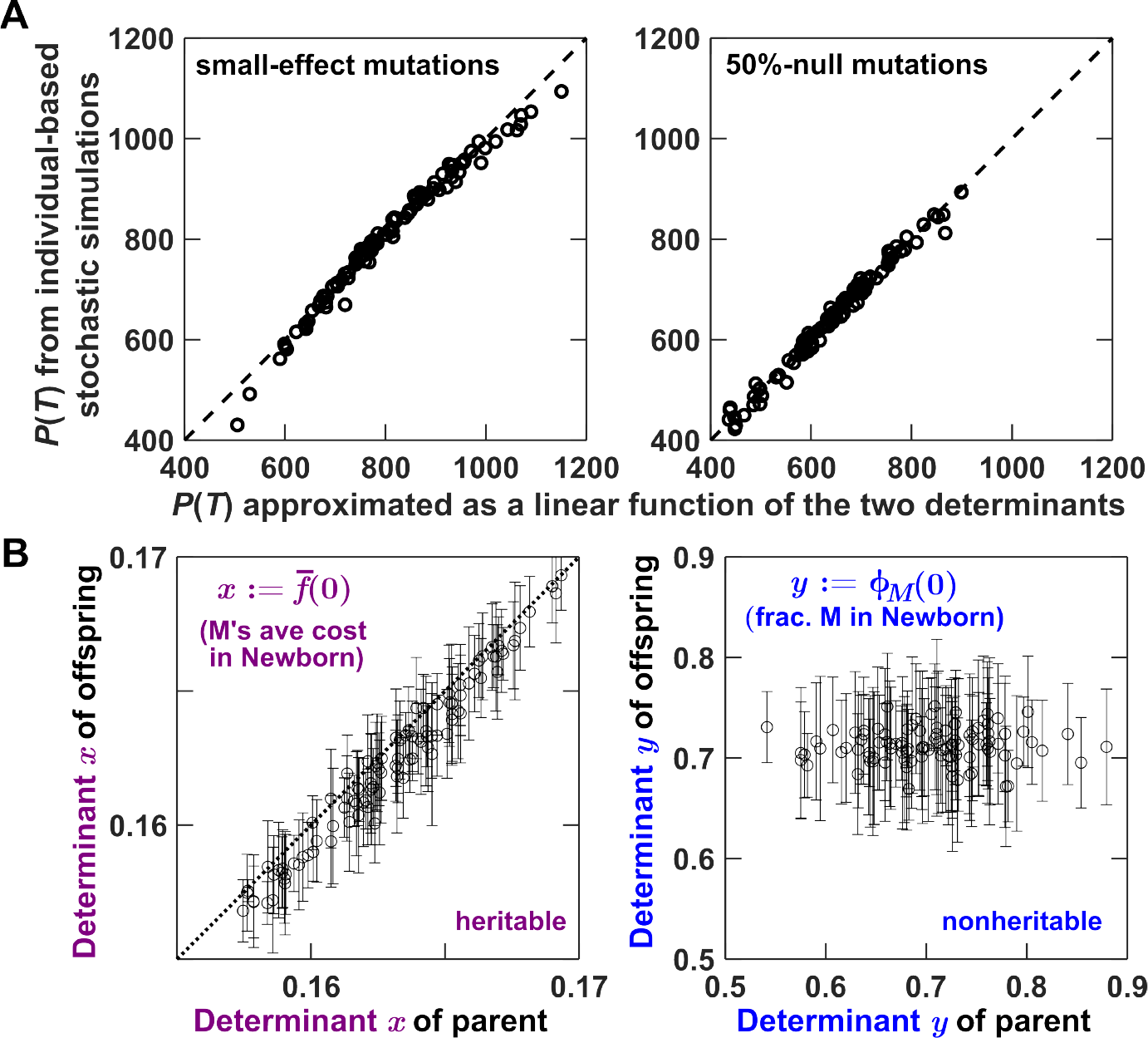
The trait of the H-M community can be approximated as a linear function of a heritable genotype determinant 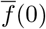 and a nonheritable species abundance determinant *ϕ*_*M*_ (0). **(A)** Under both mutation spectra (“small-effect” and “50%-null”), the true community traits *P* (*T*) obtained from individual-based stochastic simulations agree well with the linearly approximated community trait. The black dashed line is the identity line. (**B**) The genotype determinant 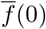 is heritable while the species abundance determinant *ϕ*_*M*_ (0) is nonheritable. The two determinants at the Newborn stage of the 100 parent communities (those plotted in the left panel of (A)) are plotted on the horizontal axis. Each parent community gives rise to 10 offspring Newborn communities, and the intra-lineage mean and standard deviation of offspring determinants are plotted as open circles and error bars on the vertical axis. The results in panel (B) were obtained under the small-effect mutation spectrum. All simulations in this figure were performed under the no-spiking condition.

### A mechanistic interpretation of heritability of a community trait in terms of heritability of trait determinants

Having decomposed a community trait into a sum of determinant contributions (Eq. 6, or more generally, Eq. 7), we now show how this linear model may be used to connect trait determinants to trait heritability

To begin, recall that the (biometric) heritability of a community trait *Z* is the slope of least squares linear regression of average offspring trait on parent trait (Eq. 3; Figure 1C). We can define the biometric heritability of determinants in the same way. For example, the biometric heritability of determinant *x* is

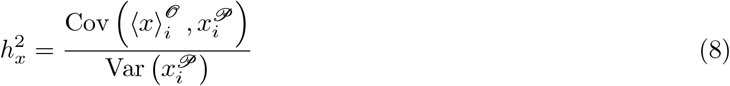

where 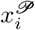 is the determinant *x* of the *i*th parent (𝒫) and 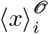 is the intra-lineage average *x* of offspring (𝒪) of the *i*th parent.

To continue, we need to make several technical assumptions. They are detailed in precise mathematical form by Eqs. S11 and S12 (in Supplement 3), but a summary is given here. Considering the linear regression of 7 applied to parents and a similar regression applied to offspring, we assume that neither residual term has a nonzero correlation with the determinant term of the other regression, and we also assume that the two residual terms are uncorrelated with each other. Additionally, considering the linear regressions that estimate offspring *x* from parent *x* or offspring *y* from parent *y*, we assume that the residual term in either such regression is uncorrelated to the parent determinant term in the other regression. With these assumptions, the biometric heritability of a community trait can be expressed in terms of the biometric heritability of the determinants *x* and *y* (Supplement Section 3):

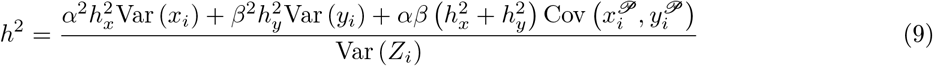

An analogous set of assumptions and an analogous version of Eq. 9 would be obtained for three or more determinants. We note that the right side of Eq. 9 can be interpreted as a community-level version of the so-called “broad-sense” heritability. In the context of individual-level selection, broad-sense heritability is the proportion of trait variance that is due to genetic variance ([26]; Supplement 2). Eq. 9 has a similar meaning because the numerator accounts for the variance and covariance of determinants weighed by their respective heritability while the denominator is the total variance of the community trait. To clarify, unless preceded by “broad-sense”, any further references to “heritability” still refer to Eq. 3 or 8.

From a technical point of view, the conditions in Eqs. S11 and S12 are sufficient to obtain Eq. 9, since it is always possible to *write down* a linear regression, regardless of whether the quantities of interest have a genuinely linear relationship; the sufficiency of Eqs. S11 and S12 may be readily verified by working through the derivation in Supplement 3. Why then did we mention Assumptions 1 and 2 of Box 2 above? This is because when the plain-language assumptions of Box 2 hold, the technical mathematical conditions of Eqs. S11 and S12 are more readily satisfied.

Let us apply this formula to the H-M community. Recall that the community trait *P* (*T*) can be approximated as a linear function of two determinants (Figure 5A; Eq. 6). The genotype determinant *x* (M’s average cost in a Newborn 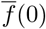 is largely heritable (Figure 5B left). This is because with sufficiently slow evolution (Assumption 1 in Box 2), if a parent community has a higher value of 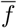 as a Newborn, then it will also tend to have a higher value of 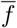 as an Adult, and so will its offspring Newborns. In contrast, the species abundance determinant *y* (fraction of M’s biomass in a Newborn *ϕ*_*M*_ (0)) is not heritable (Figure 5B right). This is because with obligatory commensal interaction [27] and our parameter choices, H and M coexist stably at a steady state value *ϕ*_*SS*_. Specifically, species compositions in parent Newborns fluctuate stochastically due to sampling errors, but converge to the steady state value *ϕ*_*SS*_ in parent Adults (which is similar across communities within a cycle due to inter-community selection; Assumption 2 in Box 2; Figure S4). Consequently, species compositions of offspring Newborns fluctuate around *ϕ*_*SS*_, uncorrelated with those of parent Newborns 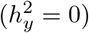. Furthermore, the genotype and the species abundance determinants are not correlated (Figure S5):

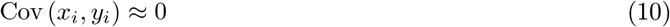

Consequently, for the H-M community, the second and third terms in the numerator of Eq. 9 are negligible. Thus, Eq. 9 becomes:

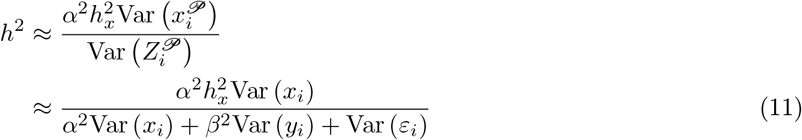

where the superscript *𝒫* is dropped in the last line (see derivation in Supplement 3). This equation indicates that increases in in the heritability of *x* or increases in the variance of *x* will tend to increase the community trait heritability. Conversely, increases in the variance of *y* (e.g. less precise pipet transfers) or in the variance of *ϵ*_*i*_ (e.g. larger random instrument errors in measurements of *P* (*T*)) will tend to decrease the community trait heritability.

### Intra-community evolution can distort the correlation between heritability and selection response by affecting all three terms in the Price equation

Higher heritability may not correlate with higher selection response because the inheritance of community traits is impacted by intra-community ecology and evolution. Inheritance is characterized by heritability (Figure 1C, olive), infidelity (Figure 1C, green), and additional terms if the parent-offspring relationship is nonlinear. For phenotypes of asexual organisms, inheritance can be well characterized by heritability due to a linear parent-offspring relationship and negligible infidelity (since mutations are rare and their average effect is small [28, 29, 30, 31, 32, 33]). In contrast, the inheritance of community traits depends on the inheritance of species abundance and genotype determinants, which can be compromised by, for example, ecological interactions and evolution during community maturation.

When intra-community evolution is slow (under small-effect mutation spectrum), biometric heritability of the community trait indeed predicts selection efficacy (i.e. community trait improves when and only when biometric heritability exceeds 0; pink regions in Figure 2(B, C)). However, when intra-community evolution is fast (under 50%-null mutation spectrum), positive heritability may not predict community trait improvement (Figure 2D). Below, we demonstrate that intra-community evolution can distort the correlation between selection response and heritability by simultaneously affecting all three terms in the Price equation (Eq. 4; Figure 1D), starting with the second term.

Second term (“infidelity”): Intra-community evolution decreases selection response for costly community traits by favoring fast-growing low-contributors. For the H-M community trait, since the residual terms 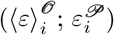 in Eq. 6 average to zero, the average trait change across one cycle in the absence of selection is:

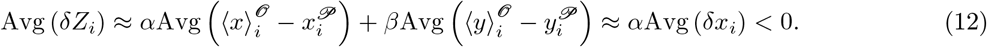

This is because 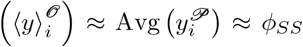 (both at the steady-state value of ∼0.7 in right panel of Figure 5B) and because intra-community evolution favors variants with low values of the cost parameter (M’s *f*) and thus the heritable determinant *x* (the average cost parameter among M cells in a Newborn) generally declines from parent Newborn to offspring Newborn 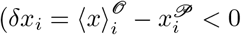, Figure 6C(i-iii)). The reduction in *x* is due to intra-community evolution, which is faster under the 50%-null mutation spectrum (Figure 6, dashed lines in C(ii) more negative than those in C(i)). The reduction is even greater when the average cost of communities are high due to effective selection (with 30%-H spiking reproduction) so that the null-mutant subpopulation of M cells increases most rapidly over maturation (dashed line in Figure 6C(iii) being the most negative).

**Figure 6:**
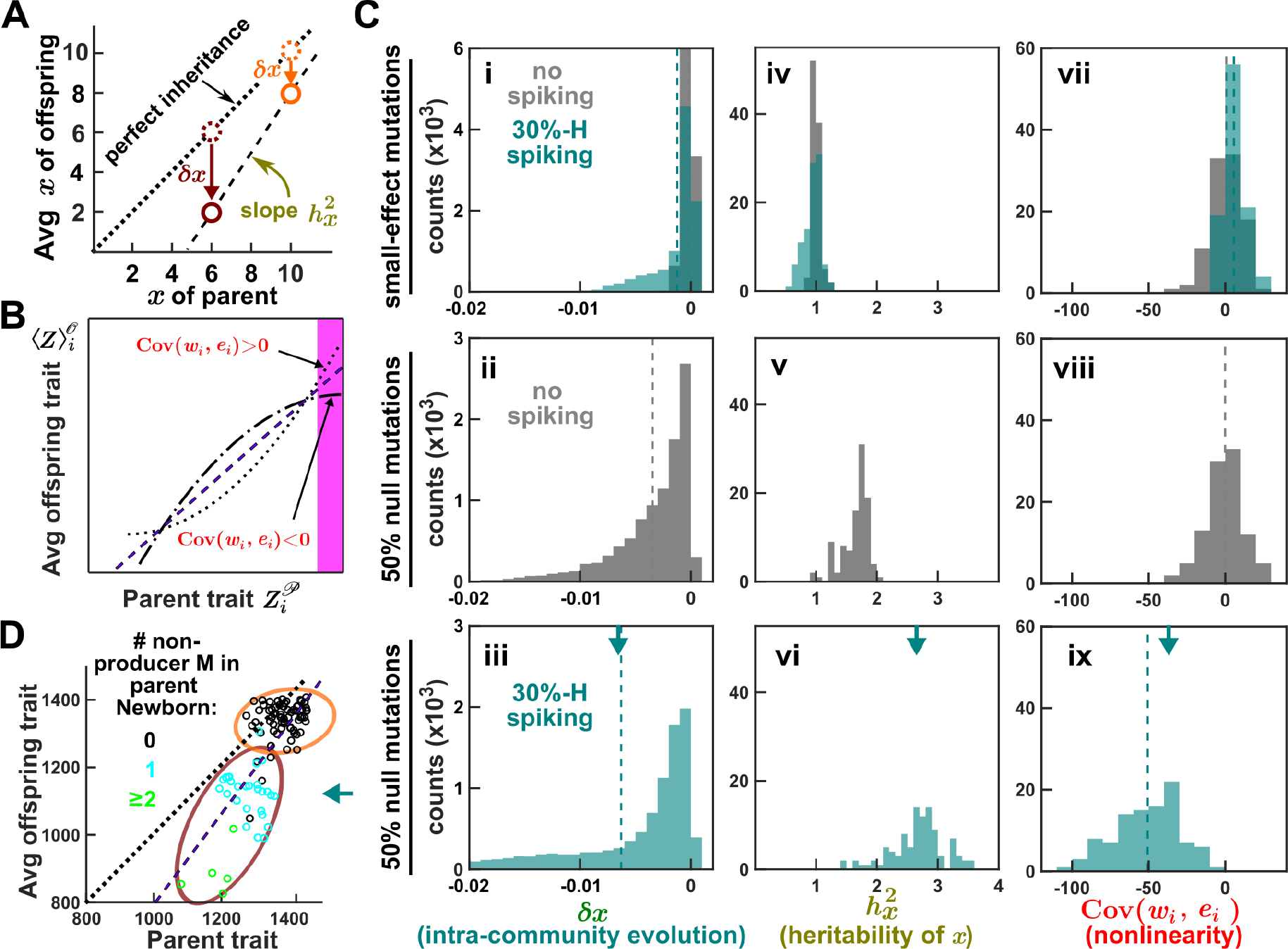
Intra-community evolution can affect all three terms in the Price equation. (**A**) Cartoon illustration of why fast intra-community evolution (e.g. under the 50%-null mutation spectrum) can inflate 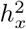 beyond 1. Newborn communities with high average cost *x* (orange circles) have zero non-producing M, and thus the decrease in *x* from parent to offspring (*6x*) is mild. In contrast, Newborn communities with low *x* (brown circles) have nonproducing M, and thus the decrease in *x* from parent to offspring is more severe as nonproducers rapidly rise in frequency. Consequently, heritability of 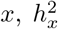 (slope of the purple dashed line), can be inflated beyond 1. (**B**) Cartoon illustration of nonlinear parent-offspring relationship leading to nonzero nonlinearity term Cov (*w*_*i*_, *e*_*i*_). When the true parent-offspring relationship is nonlinear, as depicted either by the dash-dot curve or the dotted curve, the linear regression results in the dashed line. For the dash-dot curve, residuals of the selected communities (in the magenta region) are negative, leading to a negative covariance between fitness *w* and residual *e* (Cov (*w*_*i*_, *e*_*i*_)). For the dotted curve, Cov (*w*_*i*_, *e*_*i*_) is positive. (**C**) Faster intra-community evolution leads to worse infidelity, inflated heritability, and greater nonlinearity. (**i-iii**) Intra-community evolution, characterized by the mean change in *x* from parent to offspring (Avg(*δx*) = Avg (⟨*x*⟩^𝒪^ *x*^*𝒫*^), is the distance between the vertical dashed line to zero. Intra-community evolution is slow in (i) with small-effect mutations (with or without 30%-H spiking), fast in under 50%-null mutation spectrum when the average cost in a community is low (purple in Figure 2Di). Intra-community evolution is the fastest in (iii) under the 50%-null mutation spectrum when the average cost of communities is high due to successful community selection (purple in Figure 2Ei). In this case, non-producing M has the highest growth advantage. (**iv-vi**) 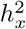 can be inflated beyond 1 under 50%-null mutation spectrum but not under small-effect mutation spectrum. (**vii-ix**) Intra-community evolution can lead to a negative nonlinearity term Cov (*w*_*i*_, *e*_*i*_). The distributions of Cov (*w*_*i*_, *e*_*i*_) center near zero except in the bottom panel where intra-community evolution is the fastest. (**D**) An example from C(iii, vi, ix) illustrates how the heritability of a community trait can exceed 1 and how a negative nonlinearity term can arise. D corresponds to Cycle 100 of Figure 2E. See the main text for details. The values of *δx*, 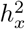 and Cov (*w*_*i*_, *e*_*i*_) from (D) are marked with teal arrows in the bottom row of (C). All histograms are computed from a total of 10^4^ communities over 100 cycles.

First term*x*(heritability selection intensity): intra-community evolution can magnify *h*^2^ (heritability of community trait, Eq. 11) by inflating 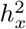 (heritability of the heritable determinant *x*). When intra-community evolution is slow (small-effect mutation), the small decline in M’s average cost (*δx*) leads to 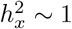 regardless of the method of reproduction (Figure 6C(iv); Eq. S21). However, when intra-community evolution is fast (50%-null mutation), the “poor get poorer” effect occurs: parent communities with lower average cost *x* tend to harbor more non-producing M whose fraction can increase significantly over maturation (Figure S2C, D), leading to a greater reduction in *x* (Figure 6A, brown compared to orange). This effect can inflate 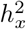 beyond 1 (Figure 6C(v, vi), Eq. S21, Figure S2C and D). By the same token, parents with low trait values generate offspring with even lower trait values (Figure 6D, blue and green circles in the brown oval farther away from the black identity line than black circles in the orange oval), leading to magnified heritability of the community trait *h*^2^ which can exceed 1 (slope of the dashed linear regression line in Figure 6D is 1.3, Figure S2C, D). Note that for a diploid individual trait with 1 locus and 2 alleles, 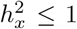 in the absence of dominance (Supplement 2). To our knowledge, trait heritability in excess of 1 has not been observed for individual phenotypes.

Third term (“nonlinearity”): Intra-community evolution can affect the nonlinearity term in the Price equation (Eq. 4):

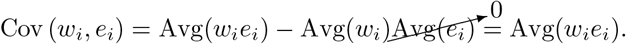

If the chosen communities have equal fitness *w*, then Cov (*w*_*i*_, *e*_*i*_)= *w* Avg(*e*_*chosen*_) and the nonlinearity term is proportional to the average residual among the chosen communities. If the dependence of a community trait on its determinants is nonlinear, the parent-offspring relationship of a community trait is in general nonlinear (Supplement Section 4) and Cov (*w*_*i*_, *e*_*i*_) may not be zero (Figure 6B). Figure 6D illustrates an example where Cov (*w*_*i*_, *e*_*i*_) is negative due to fast intra-community evolution: The communities in the cluster outlined in brown result in a large slope of the dashed regression line, which in turn results in large negative residual for the chosen communities (right tip of the upper cluster) and hence a negative covariance Cov (*w*_*i*_, *e*_*i*_) (dash-dotted line in Figure 6B, Figure 6C(ix)). In contrast, when intra-community evolution is slower, the nonlinearity term fluctuates around zero with a much smaller magnitude (Figure 6C(vii, viii)) due to an approximately linear relationship between parent and offspring traits (Figure S2(A-C)).

In sum, intra-community evolution impacts all three terms of the Price equation and can drives the previously observed correlations among them (Figures 3 and S3). Consequently, increased heritability doesn’t necessarily mean increased selection response, as shown in Figure 2.

### Strategies to increase selection efficacy

Heritability plays a pivotal role in individual selection since it is experimentally measurable and often predictive of the selection response. In contrast, we have seen that for community selection, heritability may no longer predict the selection response due to intra-community evolution. However, if heritability can be increased without a drastic impact on intra-community evolution, then the community-level selection response can still be improved.

This can be demonstrated with the H-M community trait, which has a genotype determinant 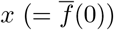and a species abundance determinant *y* (= *ϕc*_*M*_ (0)). Since the heritability of determinant *y* is nearly 0 (Figure 5B right, Figure S4), the heritability of a community trait can be approximated by Eq. 11 (top row of Table 1). Heritability can be increased, with or without accelerating intra-community evolution. For example, two methods reduce nonheritable variations in the denominator [1]: using cell sorting to reduce stochastic fluctuations in species composition Var 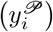 (Table 1 row a) or reducing measurement noise of the community trait Var 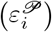 (Table 1 row c). Two other methods reduce the dependence of the community trait on nonheritable determinants (increasing *α* while decreasing *α* [21]): 30%-H spiking (Table 1 row d; Figure 2C, E) or increasing maturation time to exhaust Resource (Table 1 row e). These four methods all improve heritability without significantly accelerating intra-community evolution, and thus improve selection response [1, 21]. In contrast, since intra-community evolution is impacted by mutation rate, mutation effect, and the effective population size [34], strategies that alter these aspects (e.g. Table 1 rows b, f, and g) should be employed with caution, and evaluated in pilot studies. We summarize the mechanisms and effects of different selection strategies in Table 1.

**Table 1:**
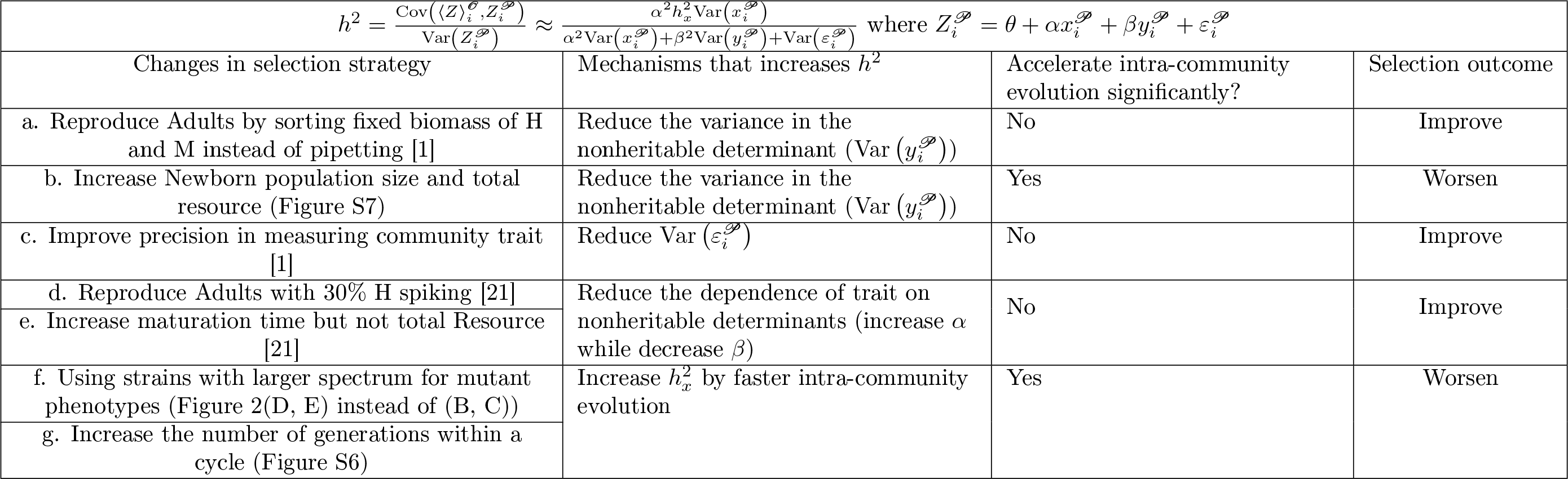
Different selection strategies increase the heritability of H-M community trait through different mechanisms but can lead to different selection outcomes depending on whether intra-community evolution is accelerated.

## Discussion

In this work, we applied the Price equation to the problem of artificial community selection. We focused on community traits that can be predicted from genotype determinants and species abundance determinants, which requires sufficiently slow intra-community evolution (Box 2, Assumption 1). Although the dependence of community traits on determinants is often nonlinear ([35]), if the inter-community distribution of determinants is narrow (Box 2, assumption 2), a linear approximation may be adequate (Figure 4; Supplement 1). The linearity assumption allowed us to interpret the heritability of a community trait (defined as the slope of parent-offspring linear regression in Eq. 3) in terms of the heritability of trait determinants (Eq. 9).

Heritability, an important component of inheritance, may [36, 6, 10, 11] or may not be predictive of selection response of communities, as observed experimentally [8, 4]. We show that intra-community evolution can affect all three terms of the Price equation and thus render heritability un-predictive of the selection response. This is in contrast to selection on individual phenotypes, where heritability is often predictive of the selection response and can be calculated from the breeder’s equation *h*^2^ = *R/S*, where *R* is the selection response (the change in the mean phenotype between generations) and *S* is the selection intensity or selection differential (the difference between the mean phenotype among parents selected for breeding and that among all parents, 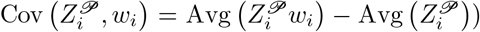. While the breeder’s equation has been applied to community selection (e.g., [37]), it might not be valid in general as higher heritability doesn’t always lead to higher selection response (Table 1). However, when heritability of a community trait is increased without drastically affecting intra-community evolution (Table 1), selection response improves (Figure 2 C vs B, E vs D).

Heritability and variation of determinants for a community trait are more complex than those for an individual phenotype. The determinants of an individual phenotype are the individual’s genotype and environment. The genotype remains nearly constant over individual maturation, and environment can be held constant in an experiment. In contrast, community trait determinants include Newborn genotype composition and species abundance. They can change drastically from parent to offspring due to 1) ecology and evolution over community maturation; and 2) stochastic fluctuations during community reproduction. In addition, intra-community variation of genotype values and inter-community variation of determinants are coupled. In the example of H-M communities, distribution of cost *f* within parent Adults determine may be related to the distribution of the determinant 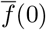 (average cost in a Newborn) among its offs<u>p</u>ring by the central limit theorem. While a large degree of inter-community variation in the determinant 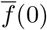 can facilitate selection, a large degree of intra-community variation in *f* can result in large decrease in 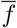 over maturation due to intra-community evolution. It is thus tricky to balance inter-community variation with undesirable intra-community evolution.

One important future direction is to understand how departure from the simplifying assumptions might affect our conclusions. For example, we assumed linear dependence of a trait on determinants, a common starting point in quantitative genetics (e.g. Ref. [23, 38]). Similar to the nonlinear terms in more complex models of individual traits (e.g., epistasis), nonlinear terms can also be included for community traits. Additionally, to arrive at the informative form of heritability in Eq. 11, we assumed no covariance between the residuals of trait-determinant regressions from different generations (Eq. S11), between a residual and a determinant of trait-determinant regressions from different generations (Eq. S11), between parent determinants and residuals of parent-offspring regressions from different traits (Eq. S12), and between different determinants within a generation (Eq. 10). While these assumptions are reasonable for the H-M community simulated in this work, they are by no means universal. For example, *x*_*i*_ and *y*_*i*_ in general are coupled (e.g. species abundance determinant *ϕ*_*M*_ (0) fluctuates around the steady state species composition *ϕ*_*SS*_ which is a function of the genotype determinant 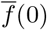; Figure S4C). Nevertheless, when the inter-community distribution of *x*_*i*_ 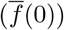 within one cycle is tight enough (Figure S4B), *y*_*i*_ (*ϕ*_*M*_ (0)) fluctuates around a nearly constant *ϕ*_*SS*_ (over the narrow red bar in Figure S4C) so that Cov (*x*_*i*_, *y*_*i*_) ≠ 0. However, when the inter-community distribution of *x*_*i*_ is wide, or when *y*_*i*_ strongly depends on *x*_*i*_, one may observe Cov (*x*_*i*_, *y*_*i*_) ≠ 0.

Another important future direction is to investigate the impact of intra-community ecology on community selection. In this work, the species abundance determinant *ϕ*_*M*_ (0) of the H-M community is nonheritable due to species composition converging to the steady-state value *ϕ*_*SS*_ during community maturation (Figure S4). In contrast, if species composition does not reach the steady state value within a cycle, then the species abundance determinant can also become heritable. While this should increase community trait heritability by increasing 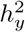 in Eq. 9, Chang et al. [11] found that selection is ineffective before communities “reach their ecological equilibrium”, or steady-state species composition. More work is needed to improve our understanding of the impact of intra-community ecology on heritability and on community selection.

In quantitative genetics, an individual phenotype is often modeled as a sum of contributions from genes, the environment, and interactions among them. Taking into consideration biological processes involved in gene transmission, theories and methods have been developed to reveal how these contributions determine a phenotype and its variations. These theories and methods, combined with records of environment and pedigrees, can help to predict selection response and thus guide the design of breeding programs. By generalizing the concept of heritability from individual phenotypes to community traits, we hope to set the stage for generalizing quantitative genetics theories and tools to guide analysis and selection on community traits. For example, with the technical advances in DNA markers and sequencing, quantitative trait locus analysis can map an individual trait to its genetic basis, such as the number and locations of responsible loci. If we allow communities to “mate” by mixing them before reproduction, can the same statistical methods be adapted? If the highly developed toolbox of quantitative genetics can be adapted to help us achieve a better understanding of a community trait’s determinants and how they are impacted by intra-community evolution and ecology, we might be able to design more effective strategies for microbial community selection.

