## Supplement for "A quantitative genetics framework for understanding the selection response of microbial communities"

#### Contents

**1 Community trait as a linear function of trait determinants 20**

**2 Individual traits: linking biometric heritability $h^2$ to broad-sense heritability $h^2_G$ 21**

**3 Community traits: linking biometric heritability $h^2$ to broad-sense heritability $h^2_G$ 24**

**4 Nonlinear dependence of a community trait on determinants leads to nonlinear parent-offspring relationship in community trait 25**

**5 H-M community: the community trait and its determinants 26**

**6 Simulating the selection of H-M communities 27**

**7 High intra-community evolution can inflate the biometric heritability of a community trait by inflating the biometric heritability of the heritable determinant 28**

### 1 Community trait as a linear function of trait determinants

In Figure 4, we illustrate how Assumption 1 (tight and nearly static distribution of genotype values within a community) and Assumption 2 (tight distribution of determinants among communities within a cycle) allow us to approximate the trait of a single-species community as a linear function of determinants. In this case, the trait is the total population size of an Adult community, while the average growth rate and the population size in a Newborn are the genotype determinant and the species abundance determinant, respectively.

Mathematically speaking, if the population is clonal and growth is deterministic and unrestricted by nutrients, population size $M(t)$ can be described by:

$$\frac{dM}{dt} = gM \tag{S1}$$

where $g$ is the growth rate. The community trait $M(T)$ (population size at time $T$) can then be solved as

$$M(T) = M(0)\exp(gT) \tag{S2}$$

Eq. S2 requires that growth rate $g$ is identical for all individuals during community maturation. In reality, $g$ can fluctuate due to noise in gene expression, or can mutate. If the distribution of $g$ within a community is narrow and does not change much over maturation (Assumption 1), then Eq. S1 is still valid if we replace $g$ with $\overline{g}(0)$ (the average $g$ in a Newborn community):

$$M(T) \approx M(0)\exp\left(\overline{g}(0)T\right) \tag{S3}$$

In other words, the community trait $M(T)$ can be predicted from two quantities: Newborn average genotype ($\overline{g}(0)$) and Newborn species abundance (Newborn population size $M(0)$). When these two quantities also vary among communities, they are determinants.

In general, a community trait depends on determinants in a nonlinear fashion (e.g. Eq. S3). However, if determinants are tightly distributed among communities (Assumption 2), then a nonlinear function can be approximated as a linear function. In this example, let the genotype determinant, species abundance determinant and community trait of the $i$th community be $\overline{g}_i(0)$, $M_i(0)$ and $M_i(T)$, respectively. Denote the inter-community average of the two determinants as $\langle\langle\overline{g}\rangle\rangle$ and $\langle\langle M\rangle\rangle$, respectively (we omitted "(0)" for simplicity). As long as $M_i(0) - \langle\langle M\rangle\rangle$ and $\overline{g}_i(0) - \langle\langle\overline{g}\rangle\rangle$ are sufficiently small, we can then linearly expand Eq. S3:

$$\begin{aligned}
M_i(T) &\approx M_i(0)\exp\left(\overline{g}_i(0)T\right) \\
&= M_i(0)\exp\left(\overline{g}_i(0)T - \langle\langle\overline{g}\rangle\rangle T + \langle\langle\overline{g}\rangle\rangle T\right) \\
&\approx \exp\left(\langle\langle\overline{g}\rangle\rangle T\right)\left(M_i(0) - \langle\langle M\rangle\rangle + \langle\langle M\rangle\rangle\right)\left[1 + (\overline{g}_i(0) - \langle\langle\overline{g}\rangle\rangle)T\right] \\
&= \exp\left(\langle\langle\overline{g}\rangle\rangle T\right)\left[(M_i(0) - \langle\langle M\rangle\rangle) + \langle\langle M\rangle\rangle + \langle\langle M\rangle\rangle(\overline{g}_i(0) - \langle\langle\overline{g}\rangle\rangle)T\right] \\
&\approx \exp\left(\langle\langle\overline{g}\rangle\rangle T\right)\left[M_i(0) + \langle\langle M\rangle\rangle(\overline{g}_i(0) - \langle\langle\overline{g}\rangle\rangle)T\right] \\
&= \exp\left(\langle\langle\overline{g}\rangle\rangle T\right)M_i(0) + \exp\left(\langle\langle\overline{g}\rangle\rangle T\right)\langle\langle M\rangle\rangle\,\overline{g}_i(0)T - \exp\left(\langle\langle\overline{g}\rangle\rangle T\right)\langle\langle M\rangle\rangle\langle\langle\overline{g}\rangle\rangle T
\end{aligned}$$

where we have used the Taylor expansion $e^x \sim 1 + x$ for $x \sim 0$ and ignored higher-order terms such as $(M_i(0) - \langle\langle M\rangle\rangle)(\overline{g}_i(0) - \langle\langle\overline{g}\rangle\rangle T)$. We can then define variables in the above equation as

$$\underbrace{M_i(T)}_{Z_i} \approx \underbrace{-\exp\left(\langle\langle\overline{g}\rangle\rangle T\right)\langle\langle M\rangle\rangle\langle\langle\overline{g}\rangle\rangle T}_{\theta} + \underbrace{\exp\left(\langle\langle\overline{g}\rangle\rangle T\right)T\langle\langle M\rangle\rangle}_{\alpha}\underbrace{\overline{g}_i(0)}_{x_i} + \underbrace{\exp\left(\langle\langle\overline{g}\rangle\rangle T\right)}_{\beta}\underbrace{M_i(0)}_{y_i}. \tag{S4}$$

This allows us to write the community trait $Z$ of the $i$th community as

$$Z_i = \theta + \alpha x_i + \beta y_i + \varepsilon_i \tag{S5}$$

where $\theta$ is a constant (intercept of the linear regression), $x_i$ is the average genotype in Newborn, $y_i$ is the species abundance in Newborn, and $\varepsilon_i$ is the residual that accounts for all the stochastic and higher order effects.

As a more complex example, consider a 3-species community whose community trait is the total population size at time $T$. Suppose that for clonal populations, $N_i$, the population size of species $i$, is described by the general Lotka-Volterra model:

$$\frac{dN_i}{dt} = r_i N_i \left(1 - \frac{\sum_{j=1}^{3} a_{ij} N_j}{K_i}\right) \tag{S6}$$

where $i,\ j \in \{1, 2, 3\}$. $r_i$ and $K_i$ represent the maximal growth rate and the carrying capacity of species $i$, respectively, while $a_{ij}N_j$ describes how species $j$ reduces the growth rate of species $i$. If species $i$ has $s_i$ strains, then the term of $N_i$ expands into $s_i$ terms ( $N_i^1, ..., N_i^{s_i}$), and parameters $r_i$, $K_i$ and $a_{ij}$ similarly expand. For example, $a_{ij}^k N_j$ represents how species $j$ (with a total population size of $N_j = \sum_{m=1}^{s_j} N_j^m$) affects strain $k$ of species $i$. The 3 equations for 3 species expand into $\sum s_i$ equations for $\sum s_i$ strains, and the equation for the population size of strain $k$ of species $i$ can be written as:

$$\frac{dN_i^k}{dt} = r_i^k N_i^k \left(1 - \frac{\sum_{j=1}^{3} a_{ij}^k \sum_{m=1}^{s_j} N_j^m}{K_i^k}\right) \tag{S7}$$

The true community trait is defined as

$$N(T) = \sum_{i=1}^{3} \sum_{k=1}^{s_i} N_i^k(T)$$

where $N_i^k(T)$ is obtained by integrating Eq. S7 from 0 to $T$. The community trait can also be predicted from species abundance and genotype determinants. There are 3 species abundance determinants: $N_i(0) = \sum_{k=1}^{s_i} N_i^k(0)$. There are 15 genotype determinants: 3 average $r_i$, 3 average $K_i$ and 9 average $a_{ij}$. Each genotype determinant is the intra-species average of the corresponding genotype, for example, the average basal growth rate of species $i$ in a Newborn is:

$$\overline{r}_i(0) = \frac{\sum_{k=1}^{s_i} N_i^k(0) r_i^k}{\sum_{k=1}^{s_i} N_i^k(0)},$$

and the average inhibition coefficient of species $j$ on species $i$ is

$$\overline{a}_{ij}(0) = \frac{\sum_{k=1}^{s_i} N_i^k(0) a_{ij}^k}{\sum_{k=1}^{s_i} N_i^k(0)}.$$

To predict community traits from these 18 determinants, replace $r_i$, $K_i$ and $a_{ij}$ in Eq. S6 with $\overline{r}_i(0)$, $\overline{K}_i(0)$ and $\overline{a}_{ij}(0)$, use the initial condition of $N_i(0) = \sum_{k=1}^{s_i} N_i^k(0)$, and then integrate over $T$. If there are $n$ communities, and if the inter-community distribution of each determinant is tight, the true community trait can be approximated as a linear function of the 18 determinants.

### 2 Individual traits: linking biometric heritability $h^2$ to broad-sense heritability $h_G^2$

Consider trait $Z$ of an asexual organism. We decompose the trait value $Z$ into the genotypic value ($G$, contribution from the genotype) and the environmental value ($E$, contribution from the environment). We assume that genotype-environment interaction ($G \times E$) is negligible. The trait of $i$th parent is thus

$$Z_i^{\mathscr{P}} = \text{Avg}(Z) + G_i^{\mathscr{P}} + E_i^{\mathscr{P}} \tag{S8}$$

where Avg($Z$) is the average population trait, and $G_i^{\mathscr{P}}$ and $E_i^{\mathscr{P}}$ represent contributions to parent trait due to deviations of the genotype and environmental values from the population averages. Offspring inherit parent's genotype value but not the environmental value ($\langle G \rangle_i^{\mathscr{O}} = G_i^{\mathscr{P}}$, $\langle E \rangle_i^{\mathscr{O}} = 0$ where $\langle \cdots \rangle$ represents intra-lineage average). The average trait for offspring of the $i$th parent is thus:

$$\langle Z\rangle_i^{\mathscr{O}} = \mathrm{Avg}(Z) + G_i^{\mathscr{P}}$$

If $\mathrm{Cov}\left(G_i^{\mathscr{P}}, E_i^{\mathscr{P}}\right) = 0$, then the biometric heritability can be estimated from Eq. 3:

$$h^2 = \frac{\mathrm{Cov}\left(\langle Z\rangle_i^{\mathscr{O}}, Z_i^{\mathscr{P}}\right)}{\mathrm{Var}\left(Z_i^{\mathscr{P}}\right)} = \frac{\mathrm{Cov}\left(G_i^{\mathscr{P}}, G_i^{\mathscr{P}} + E_i^{\mathscr{P}}\right)}{\mathrm{Var}\left(G_i^{\mathscr{P}} + E_i^{\mathscr{P}}\right)} = \frac{\mathrm{Var}\left(G_i^{\mathscr{P}}\right)}{\mathrm{Var}\left(G_i^{\mathscr{P}}\right) + \mathrm{Var}\left(E_i^{\mathscr{P}}\right)} \equiv h_G^2 \tag{S9}$$

Thus, for asexual organisms where genotypic values do not covary with environmental values, the biometric heritability is equal to the broad-sense heritability $h_G^2$ defined as the fraction of total phenotypic variation due to variation in genotypic value. For sexual organisms with recombination, if the genotypic value $G_i^{\mathscr{P}}$ depends on multiple loci, it is useful to tease out the additive effect of each locus and define the narrow-sense heritability as the fraction of total phenotypic variation due to the additive variation in genotypic value.

For asexual haploids, since $\langle G\rangle_i^{\mathscr{O}} = G_i^{\mathscr{P}}$, $h_x^2 = 1$ since $x$ in this case is $G$. For diploids, consider a genotypic value $G$ determined by a single locus with 2 alleles, A and a. The genotypic values are $G_{AA}$, $G_{Aa}$ and $G_{aa}$ for individuals with genotypes AA, Aa, and aa, respectively. Among the parent individuals who mate randomly, the frequencies of the corresponding genotypes are $f_{AA}$, $f_{Aa}$ and $f_{aa}$, respectively. The average genotypic value among parents is thus:

$$\mathrm{Avg}(G_i^{\mathscr{P}}) = f_{AA}G_{AA} + f_{Aa}G_{Aa} + f_{aa}G_{aa}$$

The frequency of alleles A and a among parents are thus $p = f_{AA} + f_{Aa}/2$ and $q = f_{aa} + f_{Aa}/2$, respectively. The average genotypic value among its offspring is thus

$$\mathrm{Avg}(\langle G\rangle_i^{\mathscr{O}}) = p^2 G_{AA} + 2pqG_{Aa} + q^2 G_{aa}.$$

The infidelity in genotypic value is then

$$\begin{aligned}
\mathrm{Avg}\left(\delta G_i\right) =& \mathrm{Avg}(\langle G\rangle_i^{\mathscr{O}}) - \mathrm{Avg}(G_i^{\mathscr{P}}) \\
=& (f_{AA}^2 + f_{Aa}^2/4 + f_{AA}f_{Aa} - f_{AA})G_{AA} + (2(f_{AA} + f_{Aa}/2)(f_{aa} + f_{Aa}/2) - f_{Aa})G_{Aa} \\
&+ (f_{aa}^2 + f_{Aa}^2/4 + f_{aa}f_{Aa} - f_{aa})G_{aa} \\
=& (f_{AA}(f_{AA} + f_{Aa} - 1) + f_{Aa}^2/4)G_{AA} + (2f_{AA}f_{aa} + f_{Aa}(f_{aa} + f_{Aa}/2 + f_{AA} - 1)G_{Aa} \\
&+ (f_{aa}(f_{aa} + f_{Aa} - 1) + f_{Aa}^2/4)G_{aa} \\
=& (-f_{AA}f_{aa} + f_{Aa}^2/4)G_{AA} + (2f_{AA}f_{aa} - f_{Aa}^2/2)G_{Aa} + (-f_{AA}f_{aa} + f_{Aa}^2/4)G_{aa} \\
=& \left(f_{AA}f_{aa} - \frac{1}{4}f_{Aa}^2\right)\left(2G_{Aa} - G_{AA} - G_{aa}\right).
\end{aligned}$$

In the absence of dominance, i.e., $G_{Aa} = \frac{1}{2}\left(G_{AA} + G_{aa}\right)$, $\mathrm{Avg}\left(\delta G_i\right) = 0$ and the infidelity term is zero.

The heritability of the genotypic value can be similarly estimated from the slope of linear regression:

$$h_x^2 = \frac{\mathrm{Cov}\left(\langle G\rangle_i^{\mathscr{O}}, G_i^{\mathscr{P}}\right)}{\mathrm{Var}\left(G_i^{\mathscr{P}}\right)}$$

where the pairs of $\left(\langle G\rangle_i^{\mathscr{O}}, G_i^{\mathscr{P}}\right)$ and the corresponding frequencies are listed in Table S1. In the absence of dominance,

$$h_x^2 = 1 - \frac{\left(G_{AA} - G_{aa}\right)^2}{\mathrm{Var}\left(G_i^{\mathscr{P}}\right)}\frac{4f_{AA}f_{aa} + f_{Aa}\left(1 - f_{Aa}\right)}{8} < 1.$$

To see this,

$$\begin{aligned}
\mathrm{Cov}\left(\langle \mathrm{G}\rangle_{\mathrm{i}}^{\mathscr{O}}, \mathrm{G}_{\mathrm{i}}^{\mathscr{P}}\right) - \mathrm{Var}(\mathrm{G}_{\mathrm{i}}^{\mathscr{P}}) &= \mathrm{E}(\langle G\rangle_i^{\mathscr{O}} G_i^{\mathscr{P}}) - \mathrm{E}(\langle G\rangle_i^{\mathscr{O}})\mathrm{E}(G_i^{\mathscr{P}}) - \mathrm{Var}(G_i^{\mathscr{P}}) \\
&= \mathrm{E}(\langle G\rangle_i^{\mathscr{O}} G_i^{\mathscr{P}}) - \mathrm{E}((G_i^{\mathscr{P}})^2) + (\mathrm{E}(G_i^{\mathscr{P}}))^2 - \mathrm{E}(\langle G\rangle_i^{\mathscr{O}})\mathrm{E}(G_i^{\mathscr{P}})
\end{aligned}$$

With no dominance, the second half is zero (since $\mathrm{E}(\langle G\rangle_i^{\mathscr{O}}) = \mathrm{E}(G_i^{\mathscr{P}})$). Thus,

$$\begin{aligned}
\mathrm{Cov}\left(\langle G\rangle_i^{\mathscr{O}}, G_i^{\mathscr{P}}\right) - \mathrm{Var}(G_i^{\mathscr{P}}) &= \mathrm{E}\left(G_i^{\mathscr{P}}\left(\langle G\rangle_i^{\mathscr{O}} - G_i^{\mathscr{P}}\right)\right) \\
&= f_{AA}G_{AA}\left(pG_{AA} + qG_{Aa} - G_{AA}\right) + f_{Aa}G_{Aa}\left(\frac{1}{2}\left(pG_{AA} + G_{Aa} + qG_{aa}\right) - G_{Aa}\right) \\
&\quad + f_{aa}G_{aa}\left(pG_{Aa} + qG_{aa} - G_{aa}\right) \\
&= \underbrace{qf_{AA}G_{AA}\left(G_{Aa} - G_{AA}\right)} + \frac{1}{2}f_{Aa}G_{Aa}\left(pG_{AA} - G_{Aa} + qG_{aa}\right) + \underbrace{pf_{aa}G_{aa}\left(G_{Aa} - G_{aa}\right)}
\end{aligned}$$

The terms marked with brackets are:

$$\begin{aligned}
& qf_{AA}G_{AA}\left(\frac{1}{2}\left(G_{AA} + G_{aa}\right) - G_{AA}\right) \\
& \quad + pf_{aa}G_{aa}\left(\frac{1}{2}\left(G_{AA} + G_{aa}\right) - G_{aa}\right) = \frac{1}{2}\left[\left(f_{aa}f_{AA}G_{AA} + f_{AA}G_{AA}f_{Aa}/2\right) - \left(f_{AA}f_{aa}G_{aa} + f_{Aa}f_{aa}G_{aa}/2\right)\right]\left(G_{aa} - G_{AA}\right) \\
& \quad = \frac{1}{2}\left[f_{aa}f_{AA}G_{AA} + f_{AA}G_{AA}f_{Aa}/2 - f_{AA}f_{aa}G_{aa} - f_{Aa}f_{aa}G_{aa}/2\right]\left(G_{aa} - G_{AA}\right) \\
& \quad = \frac{1}{2}\left[f_{aa}f_{AA}\left(G_{AA} - G_{aa}\right) + f_{AA}G_{AA}f_{Aa}/2 - f_{Aa}f_{aa}G_{aa}/2\right]\left(G_{aa} - G_{AA}\right) \\
& \quad = -\frac{f_{aa}f_{AA}}{2}\left(G_{AA} - G_{aa}\right)^2 + \frac{f_{Aa}}{4}\left(f_{AA}G_{AA} - f_{aa}G_{aa}\right)\left(G_{aa} - G_{AA}\right)
\end{aligned}$$

The middle term is:

$$\begin{aligned}
\frac{1}{2}f_{Aa}G_{Aa}\left(pG_{AA} - G_{Aa} + qG_{aa}\right) &= \frac{1}{2}f_{Aa}G_{Aa}\left(\left(f_{AA} + f_{Aa}/2\right)G_{AA} - \frac{1}{2}\left(G_{AA} + G_{aa}\right) + \left(f_{aa} + f_{Aa}/2\right)\right) \\
&= \frac{1}{4}f_{Aa}G_{Aa}\left(2f_{AA}G_{AA} + f_{Aa}G_{AA} - G_{AA} - G_{aa} + 2f_{aa}G_{aa} + f_{Aa}G_{aa}\right) \\
&= \frac{1}{4}f_{Aa}G_{Aa}\left(f_{AA}G_{AA} - f_{aa}G_{AA} + f_{aa}G_{aa} - f_{AA}G_{aa}\right) \\
&= \frac{1}{4}f_{Aa}G_{Aa}\left(f_{AA} - f_{aa}\right)\left(G_{AA} - G_{aa}\right)
\end{aligned}$$

Combining all terms:

| parent genotypic value | parent freq. | average offspring genotypic value |
|---|---|---|
| $G_{AA}$ | $f_{AA}$ | $pG_{AA}+qG_{Aa}$ |
| $G_{Aa}$ | $f_{Aa}$ | $\frac{1}{2}\left(pG_{AA}+G_{Aa}+qG_{aa}\right)$ |
| $G_{aa}$ | $f_{aa}$ | $pG_{Aa}+qG_{aa}$ |

Table S1: Average offspring genotypic value for each parent genotype.

$$\begin{aligned}
\mathrm{Cov}\left(\langle G\rangle_i^{\mathscr{O}},G_i^{\mathscr{P}}\right)-\mathrm{Var}(G_i^{\mathscr{P}}) &= -\frac{f_{aa}f_{AA}}{2}\left(G_{AA}-G_{aa}\right)^2+\frac{f_{Aa}}{4}\left(f_{AA}G_{AA}-f_{aa}G_{aa}\right)\left(G_{aa}-G_{AA}\right)\\
&\quad +\frac{1}{4}f_{Aa}G_{Aa}\left(f_{AA}-f_{aa}\right)\left(G_{AA}-G_{aa}\right)\\
&= -\frac{f_{aa}f_{AA}}{2}\left(G_{AA}-G_{aa}\right)^2+\frac{f_{Aa}}{4}[\left(f_{AA}G_{AA}-f_{aa}G_{aa}\right)\left(G_{aa}-G_{AA}\right)\\
&\quad +G_{Aa}\left(f_{AA}-f_{aa}\right)\left(G_{AA}-G_{aa}\right)]\\
&= -\frac{f_{aa}f_{AA}}{2}\left(G_{AA}-G_{aa}\right)^2+\frac{f_{Aa}}{4}\left(G_{AA}-G_{aa}\right)\big[\left(-f_{AA}G_{AA}+f_{aa}G_{aa}\right)\\
&\quad +\frac{1}{2}\left(G_{AA}+G_{aa}\right)\left(f_{AA}-f_{aa}\right)\big]\\
&= -\frac{f_{aa}f_{AA}}{2}\left(G_{AA}-G_{aa}\right)^2\\
&\quad +\frac{f_{Aa}}{8}\left(G_{AA}-G_{aa}\right)\left[-f_{AA}G_{AA}+f_{aa}G_{aa}+G_{aa}f_{AA}-G_{AA}f_{aa}\right]\\
&= -\frac{f_{aa}f_{AA}}{2}\left(G_{AA}-G_{aa}\right)^2+\frac{f_{Aa}}{8}\left(G_{AA}-G_{aa}\right)\left[+f_{aa}(G_{aa}-G_{AA})+(G_{aa}-G_{AA})f_{AA}\right]\\
&= -\frac{f_{aa}f_{AA}}{2}\left(G_{AA}-G_{aa}\right)^2-\frac{f_{Aa}}{8}\left(G_{AA}-G_{aa}\right)^2\left(f_{aa}+f_{AA}\right)\\
&= -\left(G_{AA}-G_{aa}\right)^2\left[\frac{f_{aa}f_{AA}}{2}+\frac{f_{Aa}}{8}\left(f_{aa}+f_{AA}\right)\right]
\end{aligned}$$

Thus,

$$\begin{aligned}
h_x^2 &= \frac{\mathrm{Cov}\left(\langle G\rangle_i^{\mathscr{O}},G_i^{\mathscr{P}}\right)}{\mathrm{Var}\left(G_i^{\mathscr{P}}\right)}\\
&= 1-\frac{\left(G_{AA}-G_{aa}\right)^2\left[4f_{aa}f_{AA}+f_{Aa}\left(f_{aa}+f_{AA}\right)\right]}{8\mathrm{Var}\left(G_i^{\mathscr{P}}\right)}
\end{aligned}$$

### 3 Community traits: linking biometric heritability $h^2$ to broad-sense heritability $h_G^2$

Begin by writing $Z$ in linear regression form, where the predictors are two determinants, $x$ and $y$. Equation 7 for parent communities then becomes:

$$Z_i^{\mathscr{P}}=\theta+\alpha x_i^{\mathscr{P}}+\beta y_i^{\mathscr{P}}+\varepsilon_i^{\mathscr{P}} \tag{S10}$$

where slopes $\alpha$ and $\beta$ indicate how strongly the community trait depends on the two determinants, $\theta$ is a constant, and $\varepsilon_i^{\mathscr{P}}$ is the residual of the linear approximation (which can include stochastic effects and any nonlinear dependence of $Z_i^{\mathscr{P}}$ on the determinants). Similarly, the trait of the $j$th offspring of the $i$th parent, $Z_{ij}^{\mathscr{O}}$, can be written as

$$Z_{ij}^{\mathscr{O}}=\theta+\alpha x_{ij}^{\mathscr{O}}+\beta y_{ij}^{\mathscr{O}}+\varepsilon_{ij}^{\mathscr{O}}$$

If we average the traits and determinants among all offspring from the same ($i$th) parent, we have

$$\langle Z\rangle_i^{\mathscr{O}} = \theta + \alpha\,\langle x\rangle_i^{\mathscr{O}} + \beta\,\langle y\rangle_i^{\mathscr{O}} + \langle\varepsilon\rangle_i^{\mathscr{O}}$$

where the subscript $j$ has been averaged out.

To write the biometric heritability of community trait $h^2$ in a simple form, we assume that the covariance between the residual terms and other terms are zero:

$$\mathrm{Cov}\left(x_i^{\mathscr{P}}, \langle\varepsilon\rangle_i^{\mathscr{O}}\right) = \mathrm{Cov}\left(y_i^{\mathscr{P}}, \langle\varepsilon\rangle_i^{\mathscr{O}}\right) = \mathrm{Cov}\left(\varepsilon_i^{\mathscr{P}}, \langle\varepsilon\rangle_i^{\mathscr{O}}\right) = \mathrm{Cov}\left(\varepsilon_i^{\mathscr{P}}, \langle x\rangle_i^{\mathscr{O}}\right) = \mathrm{Cov}\left(\varepsilon_i^{\mathscr{P}}, \langle y\rangle_i^{\mathscr{O}}\right) = 0 \tag{S11}$$

We then have

$$\begin{aligned} h^2 &= \frac{\mathrm{Cov}\left(\langle Z\rangle_i^{\mathscr{O}}, Z_i^{\mathscr{P}}\right)}{\mathrm{Var}\left(Z_i^{\mathscr{P}}\right)} \\ &= \frac{\mathrm{Cov}\left(\alpha\,\langle x\rangle_i^{\mathscr{O}} + \beta\,\langle y\rangle_i^{\mathscr{O}} + \langle\varepsilon\rangle_i^{\mathscr{O}}, \alpha x_i^{\mathscr{P}} + \beta y_i^{\mathscr{P}} + \varepsilon_i^{\mathscr{P}}\right)}{\mathrm{Var}\left(Z_i^{\mathscr{P}}\right)} \\ &= \frac{\alpha^2\mathrm{Cov}\left(x_i^{\mathscr{P}}, \langle x\rangle_i^{\mathscr{O}}\right) + \beta^2\mathrm{Cov}\left(y_i^{\mathscr{P}}, \langle y\rangle_i^{\mathscr{O}}\right) + \alpha\beta\left(\mathrm{Cov}\left(x_i^{\mathscr{P}}, \langle y\rangle_i^{\mathscr{O}}\right) + \mathrm{Cov}\left(y_i^{\mathscr{P}}, \langle x\rangle_i^{\mathscr{O}}\right)\right)}{\mathrm{Var}\left(Z_i^{\mathscr{P}}\right)}. \end{aligned}$$

Similar to Eq. 2, we can regress $\langle x\rangle_i^{\mathscr{O}}$ and $\langle y\rangle_i^{\mathscr{O}}$ against $x_i^{\mathscr{P}}$ and $y_i^{\mathscr{P}}$, respectively:

$$\langle x\rangle_i^{\mathscr{O}} = h_x^2 x_i^{\mathscr{P}} + \theta_x + e_{xi}$$

$$\langle y\rangle_i^{\mathscr{O}} = h_y^2 y_i^{\mathscr{P}} + \theta_y + e_{yi}$$

If we also assume that

$$\mathrm{Cov}\left(x_i^{\mathscr{P}}, e_{yi}\right) = \mathrm{Cov}\left(y_i^{\mathscr{P}}, e_{xi}\right) = 0, \tag{S12}$$

then we have

$$\begin{aligned} h^2 &= \frac{\mathrm{Cov}\left(\langle Z\rangle_i^{\mathscr{O}}, Z_i^{\mathscr{P}}\right)}{\mathrm{Var}\left(Z_i^{\mathscr{P}}\right)} \\ &= \frac{\alpha^2 h_x^2 \mathrm{Var}\left(x_i^{\mathscr{P}}\right) + \beta^2 h_y^2 \mathrm{Var}\left(y_i^{\mathscr{P}}\right) + \alpha\beta\left(h_x^2 + h_y^2\right)\mathrm{Cov}\left(x_i^{\mathscr{P}}, y_i^{\mathscr{P}}\right)}{\mathrm{Var}\left(Z_i^{\mathscr{P}}\right)}. \end{aligned} \tag{S13}$$

Note that Eq. S13 also requires the conditions $\mathrm{Cov}\left(x_i^{\mathscr{P}}, e_{xi}\right) = 0$ and $\mathrm{Cov}\left(y_i^{\mathscr{P}}, e_{yi}\right) = 0$, but these are consequences of a basic property of simple least-squares linear regression that the residual is uncorrelated to the predictor variable. That is, having defined $h_x^2 = \mathrm{Cov}\left(\langle x\rangle_i^{\mathscr{O}}, x_i^{\mathscr{P}}\right)/\mathrm{Var}\left(x_i^{\mathscr{P}}\right)$ and $\theta_x = \mathrm{Avg}(\langle x\rangle_i^{\mathscr{O}}) - h_x^2\mathrm{Avg}(x_i^{\mathscr{P}})$ as per usual, it can be shown via algebra that $\mathrm{Cov}\left(x_i^{\mathscr{P}}, e_{xi}\right) = 0$. Equation S13 says that the heritability of a trait is influenced by the variation and heritability of each determinant, as well as the interaction among different determinants. Note that superscripts in the last row of the above equation are all $\mathscr{P}$ and thus can be dropped, yielding Eq. 9.

### 4 Nonlinear dependence of a community trait on determinants leads to nonlinear parent-offspring relationship in community trait

Consider a simple community trait $Z$ with one determinant $x$ so that the trait of a parent is $Z_i^{\mathscr{P}} = x_i^{\mathscr{P}}$. If the determinant of offspring is $x_i^{\mathscr{O}} = hx_i^{\mathscr{P}}$ where $h$ corresponds to $h_x^2$ in Eq. 8, then the traits of parent and offspring communities are linearly related since $Z_i^{\mathscr{O}} = x_i^{\mathscr{O}} = hx_i^{\mathscr{P}} = hZ_i^{\mathscr{P}}$.

In contrast, when a community trait is a nonlinear (e.g. a quadratic) function of its determinant:

$$Z_i^{\mathscr{P}} = x_i^{\mathscr{P}} + a\left(x_i^{\mathscr{P}}\right)^2$$

where $a \ll 1$, then

$$Z_i^{\mathscr{O}} = x_i^{\mathscr{O}} + a\left(x_i^{\mathscr{O}}\right)^2 = hx_i^{\mathscr{P}} + a\left(hx_i^{\mathscr{P}}\right)^2.$$

When regressing $Z_i^{\mathscr{O}}$ against $Z_i^{\mathscr{P}}$, the slope is, to the first order of $a$:

$$\begin{aligned}
\frac{\mathrm{Cov}\left(Z_i^{\mathscr{P}}, Z_i^{\mathscr{O}}\right)}{\mathrm{Var}\left(Z_i^{\mathscr{P}}\right)} &= \frac{\mathrm{Cov}\left(x_i^{\mathscr{P}} + a\left(x_i^{\mathscr{P}}\right)^2, hx_i^{\mathscr{P}} + ah^2\left(x_i^{\mathscr{P}}\right)^2\right)}{\mathrm{Var}\left(x_i^{\mathscr{P}} + a\left(x_i^{\mathscr{P}}\right)^2\right)} \\
&\approx \frac{h\mathrm{Var}\left(x_i^{\mathscr{P}}\right) + ah(1+h)\mathrm{Cov}\left(x_i^{\mathscr{P}}, \left(x_i^{\mathscr{P}}\right)^2\right)}{\mathrm{Var}\left(x_i^{\mathscr{P}}\right) + a\mathrm{Cov}\left(x_i^{\mathscr{P}}, \left(x_i^{\mathscr{P}}\right)^2\right)} \\
&= h\frac{1 + a\left(1+h\right)\mathrm{Cov}\left(x_i^{\mathscr{P}}, \left(x_i^{\mathscr{P}}\right)^2\right)/\mathrm{Var}\left(x_i^{\mathscr{P}}\right)}{1 + a\mathrm{Cov}\left(x_i^{\mathscr{P}}, \left(x_i^{\mathscr{P}}\right)^2\right)/\mathrm{Var}\left(x_i^{\mathscr{P}}\right)} \\
&\approx h(1+ahs)
\end{aligned}$$

where $s := \mathrm{Cov}\left(x_i^{\mathscr{P}}, \left(x_i^{\mathscr{P}}\right)^2\right)/\mathrm{Var}\left(x_i^{\mathscr{P}}\right)$. The parent-offspring regression is thus,

$$\begin{aligned}
Z_i^{\mathscr{O}} &= h(1+ahs)Z_i^{\mathscr{P}} + c + \varepsilon_i \\
&= h(1+ahs)(x_i^{\mathscr{P}} + a\left(x_i^{\mathscr{P}}\right)^2) + c + \varepsilon_i
\end{aligned}$$

where $c$ is the intercept and $\varepsilon_i$ is the residual. Hence, to the first order of $a$, the residual is

$$\begin{aligned}
\varepsilon_i &= hx_i^{\mathscr{P}} + ah^2\left(x_i^{\mathscr{P}}\right)^2 - h(1+ahs)(x_i^{\mathscr{P}} + a\left(x_i^{\mathscr{P}}\right)^2) - c \\
&= ah(h-1)\left(x_i^{\mathscr{P}}\right)^2 - ah^2 s x_i^{\mathscr{P}} - c
\end{aligned}$$

The dependence of $\varepsilon_i$ on $x_i^{\mathscr{P}}$ means that the parent-offspring regression on community trait is not linear.

### 5 H-M community: the community trait and its determinants

Assume that all cells have the same genotype values, that death and birth processes are deterministic, and that there is no evolution. community trait $P(T)$ can then be numerically integrated from the following set of scaled differential equations [1]:

$$\frac{dR}{dt} = -c_{RM}g_M(R, B)M - c_{RH}g_H(R)H \tag{S14}$$

$$\frac{dB}{dt} = g_H(R)H - c_{BM}g_M(R, B)M \tag{S15}$$

$$\frac{dP}{dt} = fg_M(R, B)M \tag{S16}$$

$$\frac{dH}{dt} = g_H(R)H - \delta_H H \tag{S17}$$

$$\frac{dM}{dt} = g_M(R, B)\left(1-f\right)M - \delta_M M \tag{S18}$$

where

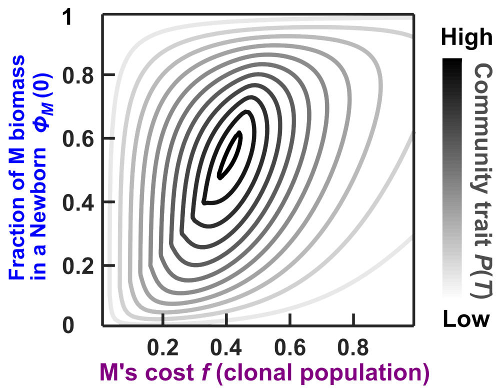


Figure S1: The community trait $P(T)$ is a nonlinear function of $\phi_M(0)$ and $f$. The contour plot is calculated by numerically integrating Eqs S14-S20 assuming that both H and M are clonal and that the total biomass in a Newborn is 100. When Assumption 1 in Box 2 is satisfied, M's cost $f$ can be replaced by the average M's cost in a Newborn, $\overline{f}(0)$.

$$g_H(R) = g_{Hmax}\frac{R}{R+K_{HR}} \tag{S19}$$

$$g_M(R,\,B) = g_{Mmax}\frac{R_M B_M}{R_M+B_M}\left(\frac{1}{R_M+1}+\frac{1}{B_M+1}\right) \tag{S20}$$

and $R_M = R/K_{MR}$ and $B_M = B/K_{MB}$.

Eq. S14 states that Resource $R$ is depleted by biomass growth of M and H, where $c_{RM}$ and $c_{RH}$ represent the amount of $R$ consumed per unit of M and H biomass, respectively. Eq. S15 states that Byproduct B is released as H grows, and is decreased by biomass growth of M due to consumption ($c_{BM}$ represents the amount of B consumed per unit of M biomass). Eq. S16 states that Product P is produced as $f$ fraction of potential M growth. Eq.S17 states that H biomass increases at a rate dependent on Resource $R$ in a Monod fashion (Eq. S19) and decreases at the death rate $\delta_H$. Eq. S18 states that M biomass increases at a rate dependent on Resource $R$ and Byproduct $B$ (the Mankad and Bungay model in Eq. S20 [39]) discounted by $(1-f)$ due to the fitness cost of making Product, and decreases at the death rate $\delta_M$. In the Monod growth model (Eq. S19), $g_{Hmax}$ is the maximal growth rate of H and $K_{HR}$ is the $R$ at which $g_{Hmax}/2$ is achieved. In the Mankad and Bungay model (Eq. S20), $K_{MR}$ is the $R$ at which $g_{Mmax}/2$ is achieved when $B$ is in excess; $K_{MB}$ is the $B$ at which $g_{Mmax}/2$ is achieved when $R$ is in excess.

If M and H are clonal and growth is deterministic, community trait $P(T)$ can be obtained by numerically integrating Eqs S14-S20. In this case, $P(T)$ depends on the initial conditions (e.g. the abundances of M and H in the Newborn) and parameters (e.g. M's cost $f$; M and H's affinity to metabolites). The dependence is nonlinear, as shown by the contour plot Figure S1.

### 6 Simulating the selection of H-M communities

In this work, the only mutable genotype is M's $f$. The other genotypes ($g_{Mmax}$, $g_{Hmax}$, $K_{MR}$ etc.) are held constant. Under the small-effect mutation spectrum, a mutation increases or decreases $f$ by on average 0.01. Under the 50%-null mutation spectrum, 50% of the time the mutation reduces $f$ to 0, creating a non-produceing M. The other 50% of the time the mutation is small-effect. The community trait is defined as the total amount of Product in an Adult community: $P(T)$.

When we simulate artificial community selection on such H-M communities, each cycle starts with a total of 100 Newborn communities. The population sizes are measured in biomass unit. The total biomass of each Newborn community, $BM_{target}$, is closely fixed to $BM_{target} = 100$ units, equivalent to 50-100 cells each having a biomass between 1 and 2 units. During a fixed maturation period $T$, the cells take up and release metabolites, grow in biomass, and divide when their biomass reach a critical value of 2. For M cells

| Parameters for phenotypes of H and M used in the simulations | | |
|---|---|---|
| | Definition | Value or range |
| $f$ | fraction of M growth diverted to producing P | 0-1 |
| $K_{MR}$ | fold of $R(0)$ at which $g_{Mmax}/2$ is achieved in excess B | 1/3 |
| $K_{MB}$ | amount of Byproduct at which $g_{Mmax}/2$ is achieved in excess R | $\frac{1}{3} \times 10^2$ |
| $K_{HR}$ | fold of $R(0)$ at which $g_{Hmax}/2$ is achieved | 1/5 |
| $g_{Mmax}$ | maximal biomass growth rate of M | 0.7 per unit time |
| $g_{Hmax}$ | maximal biomass growth rate of H | 0.3 per unit time |
| $\delta_M$ | death rate of M | $3.5 \times 10^{-3}$per unit time |
| $\delta_H$ | death rate of H | $1.5 \times 10^{-3}$ per unit time |
| $c_{RM}$ | fraction of $R(0)$ consumed per M biomass grown | $10^{-4}$ |
| $c_{RH}$ | fraction of $R(0)$ consumed per H biomass grown | $10^{-4}$ |
| $c_{BM}$ | amount of Byproduct consumed per M biomass grown | $\frac{1}{3}$ |
| $P_{\text{mut}}$ | mutation probability per cell division for each mutable phenotype | $2 \times 10^{-3}$ |

Table S2: Parameters for phenotypes of H and M used in the simulations. Among these phenotypes, only $f$ can be modified by mutations. For parameter justifications, see Ref. [1].

after each division, both daughter cells mutate with a probability of $P_{mut} = 2 \times 10^{-3}$. At the end of the maturation cycle $T$, we choose 10 Adult communities with the highest traits (the total amount of Product in an Adult $P(T)$) and allow each to reproduce 10 Newborn communities for the next cycle. Reproduction is performed by randomly assigning H and M cells to each Newborn until the total biomass in a Newborn reaches very closely to 100 units. As a result, although the total biomass in a Newborn is nearly a constant ($BM_{target} = 100$ units), the fraction of M biomass in a Newborn, $\phi_M(0)$, fluctuates randomly with a relative standard deviation of ~10%-30%.

### 7 High intra-community evolution can inflate the biometric heritability of a community trait by inflating the biometric heritability of the heritable determinant

In general

$$h_x^2 = \frac{\text{Cov}\left(\langle x\rangle_i^{\mathscr{O}}, x_i^{\mathscr{P}}\right)}{\text{Var}\left(x_i^{\mathscr{P}}\right)} = 1 + \frac{\text{Cov}\left(\delta x_i, x_i^{\mathscr{P}}\right)}{\text{Var}\left(x_i^{\mathscr{P}}\right)} \tag{S21}$$

where $\delta x_i = \langle x\rangle_i^{\mathscr{O}} - x_i^{\mathscr{P}}$ measures intra-community evolution. For the H-M community, $x$ is the average cost paid by M in a Newborn community, $\overline{f}(0)$. Since M cells with lower cost $f$ grow faster than those with higher cost $f$, the average cost of a community decreases during maturation. Consequently, offspring Newborns in general have lower average cost compared to parent Newborns. In other words, $\delta x_i = \langle x\rangle_i^{\mathscr{O}} - x_i^{\mathscr{P}} < 0$ due to the intra-community evolution in general (y axis of data points in Figure S2(A_D)(i) mostly negative). This is in contrast with individual traits of asexual individuals, where offspring usually share the same heritable determinant (genotypic value) $x$ as their parent, so that $\delta x_i = \langle x\rangle_i^{\mathscr{O}} - x_i^{\mathscr{P}} \approx 0$. Even if mutation alters $x_i^{\mathscr{P}}$, $\text{Cov}\left(x_i^{\mathscr{P}}, \delta x_i\right)$ is often 0 if random mutations lead to zero average effects.

For H-M communities, under the small-effect mutation spectrum, $\delta x_i$ is small (Figure 6C(i), Figure S2(A-B)(i)). Cov $\left(x_i^{\mathscr{P}}, \delta x_i\right)$ is then small compared to Var $\left(x_i^{\mathscr{P}}\right)$, meaning that $h_x^2 = 1 + \text{Cov}\left(x_i^{\mathscr{P}}, \delta x_i\right) / \text{Var}\left(x_i^{\mathscr{P}}\right)$ is close to 1, as shown in Figure 6C(iv). In contrast, under the 50%-null mutation spectrum, intra-community evolution is faster. This leads to more negative $\delta x_i$ and $h_x^2 > 1$ as shown in Figure S2(C-D)(i) and Figure 6C(ii-iii, v-vi). This is because in this case, $\delta x_i$ is largely driven by non-producing Ms that have a significant fitness advantage (colored circles in Figure S2C-D)(i)). If a parent community has low $\overline{f}(0)$, it's likely to have more non-producing M with $f = 0$. Its offspring communities are more likely to inherit more non-producing M that further lower $\overline{f}(0)$. Therefore, under the 50%-null mutation spectrum, $h_x^2$ is inflated due to intra-community evolution. This in turn leads to an increase in biometric heritability (Figure 2B vs. D, C vs. E; Figure S2 A(iii) vs. C(iii), B(iii) vs. D(iii)). However, the increased biometric heritability doesn't lead to increased selection response, as the second and third terms of the Price equation 4 can become more negative due to intra-community evolution.

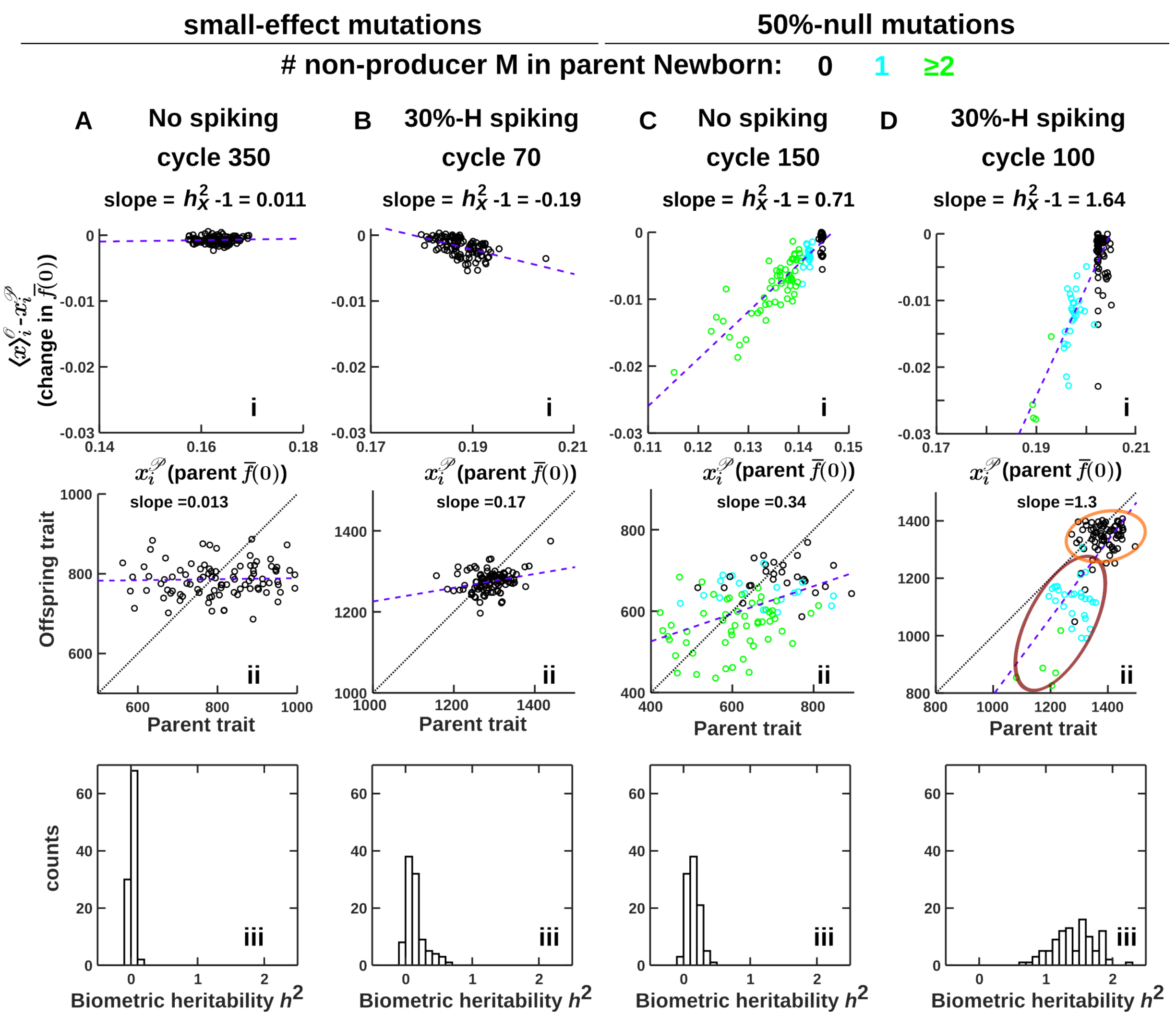


Figure S2: **Intra-community evolution can inflate heritability of a trait and its determinant.** (i) Change in determinant $x$ (M's average cost in Newborn $\overline{f}(0)$) over one cycle $\delta x_i = \langle x\rangle_i^{\mathscr{O}} - x_i^{\mathscr{P}}$ are plotted against parent determinant $x_i^{\mathscr{P}}$ for 100 H-M communities under different reproduction schemes and mutation spectra, where the purple dashed lines indicate the least square linear fitting. Compared to (A, B), decrease of $x$ in (C, D) is much greater in magnitude. In (C, D), the number of non-producing M cells ($f = 0$) at the Newborn stage is color coded. The lower the parent $x$, the greater the decline in $x$ from parent to offspring ("the poor get poorer"). (ii) 100 pairs of parent and offspring traits are scatter plotted for one cycle. The purple dashed lines indicate the linear least square regression line so the slope is the biometric heritability. D(ii) is identical to Figure 6D. (iii) The histograms of biometric heritability over 100 cycles. Under the same reproduction schemes (no-spiking or 30%-H spiking), heritability is generally higher when non-producing M speeds up intra-community evolution (compare A(iii) v.s. C(iii), B(iii) v.s. D(iii)).

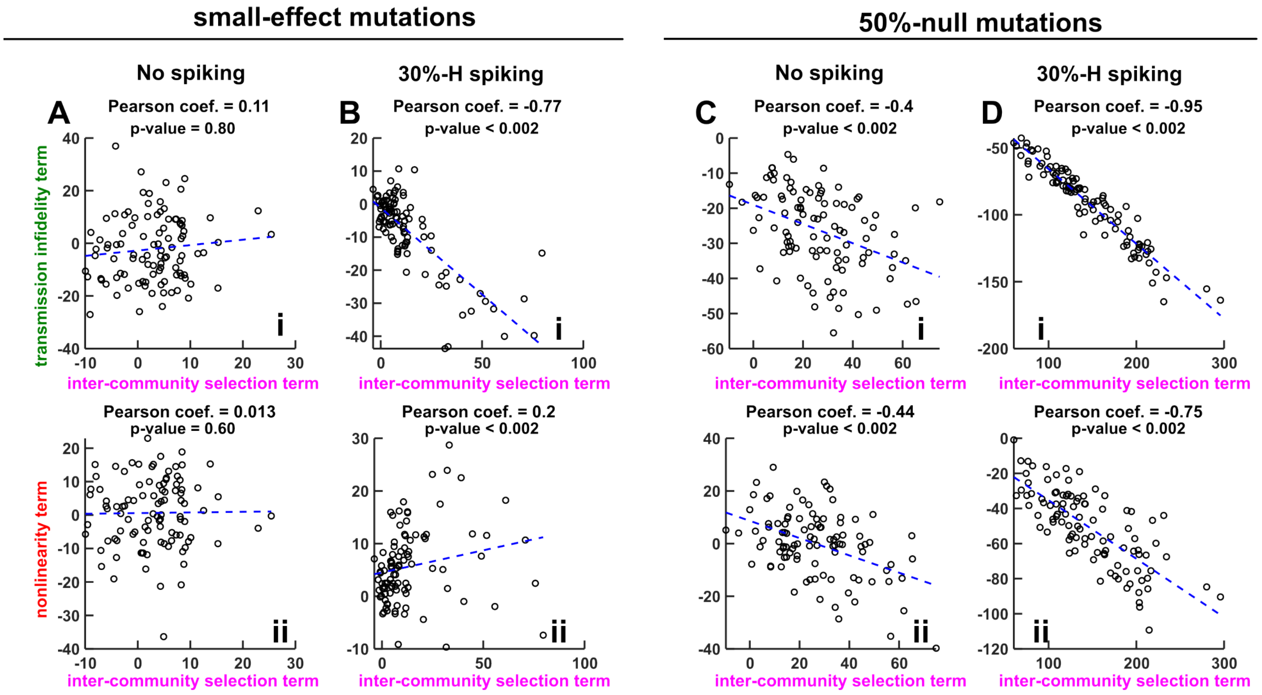


Figure S3: **Inter-community selection may be coupled to transmission infidelity and nonlinearity.** Compared to correlations under slow intra-community evolution (A, B), correlations are stronger when intra-community evolution is faster (C, D). To see whether the correlation was strong enough to indicate statistical dependence between the variables shown on the horizontal and vertical axes, we performed a nonparametric permutation hypothesis test for each panel. Specifically, under each simulation condition, we ran 9 additional simulation replicates over the same 100 cycles (for a total of 10 replicates) and performed a permutation test [40]. The test statistic is computed by calculating the Pearson correlation coefficient within each replicate, and then taking the average correlation among all 10 replicates. In principle, there are 10! different ways to permute, and thus an exact test would use 10! different permutations. However, for the sake of computational efficiency we used 1000 randomly selected permutations. To calculate the p-value, we computed the proportion of 1000 randomized data sets with a lower average correlation coefficient than the non-randomized data ($p_{lower}$), the proportion of randomized data sets with a higher average correlation coefficient than non-randomized data ($p_{higher}$), and reported the p-value as $p = 2\min(p_{lower}, p_{higher})$, where the factor of 2 corrects for using two tails. Except in the case of small-effect mutations with no spiking (A), the permutation test yielded strong evidence that the inter-community selection term is correlated with the other two terms in the Price equation.

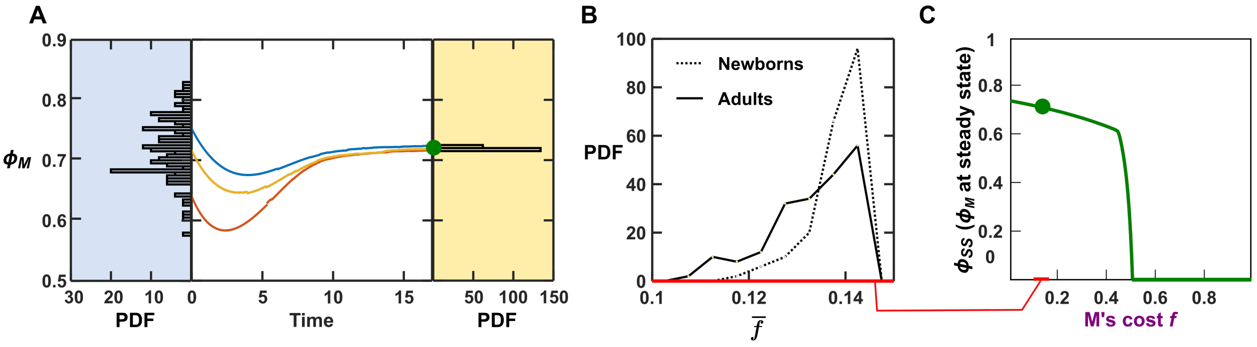


Figure S4: **The fraction of M in a community converges toward a steady state value.** (A) In Cycle 150 of the simulation shown in Figure 2D(ii), PDF (probability density function) of $\phi_M(0)$, the species abundance determinant, is shown in the left histogram with blue background. $\phi_M(t)$ of 3 communities over the maturation time are then plotted in red, yellow and blue curves, respectively. All 3 curves approaches a common steady state shown by the green dot regardless of their $\phi_M(0)$. As a result, the distribution of $\phi_M(T)$ (the fraction of M in the Adult communities) plotted on the right with a yellow background is much tighter. In this example, the inter-community averages of $\phi_M(0)$ and $\phi_M(T)$ are very similar, both being 0.72, while the inter-community standard deviation is $5.3 \times 10^{-2}$ for $\phi_M(0)$ to $2.3 \times 10^{-3}$ for $\phi_M(T)$. (B) PDF of $\overline{f}(0)$ (dotted line) and $\overline{f}(T)$ (solid line) among the same 100 communities. The PDFs are narrow as the C.V (coefficient of variation; standard deviation divided by mean) of the distributions of $\overline{f}(0)$ and $\overline{f}(T)$ are 4.2% and 7.5%, respectively. (C) The steady state of the fraction of M in an Adult, $\phi_{SS}$, depends on M's cost $f$. To calculate the species composition attractor, we integrated Eq. S14-S18 to obtain $\phi_M(T) - \phi_M(0)$ for each grid point on the 2D mesh of $\phi_M(0)$ and $f$. The contour of $\phi_M(T) - \phi_M(0) = 0$ is then the species composition attractor. Under Assumption 1, the calculation is still valid if $f$ is replaced by $\overline{f}(0)$. Even though $\phi_{SS}$ (and therefore $\phi_M(T)$) depends on $\overline{f}(0)$, when the inter-community distribution of $\overline{f}(0)$ within a cycle is narrow (the $x$ axis of B corresponds to the red bar in C), $\phi_M(T)$ among Adults is nearly constant (compare histograms on the left and right in A).

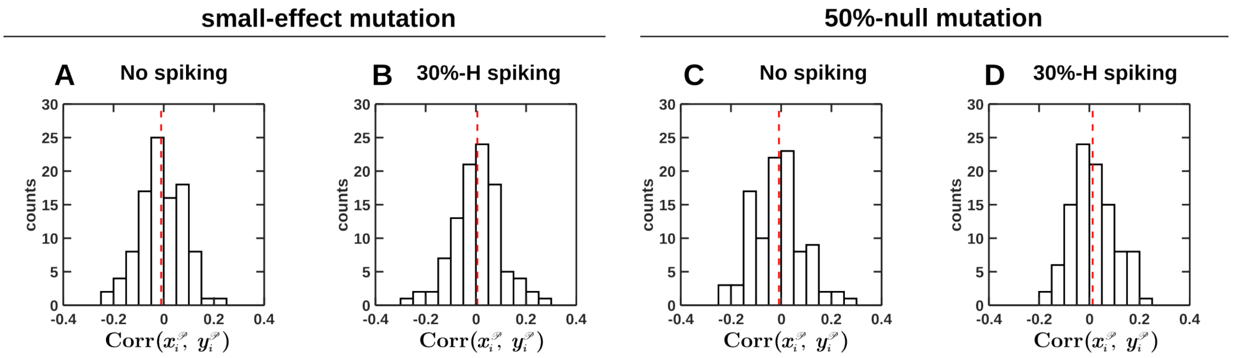


Figure S5: **The Pearson correlation coefficient between determinants $x_i^{\mathscr{P}}$ ($\overline{f}(0)$ of a community) and $y_i^{\mathscr{P}}$ ($\phi_M(0)$ of a community) is very small.** The histograms of 100 Pearson correlation coefficients Corr $\left(x_i^{\mathscr{P}}, y_i^{\mathscr{P}}\right)$ for 100 cycles are plotted with the sample mean marked by the red dashed line. In all cases, the sample mean (red dashed line) falls within the 95% confidence interval for the mean, which is calculated using Matlab's bootci function. The confidence intervals are A: [-0.03, 0.01] B: [-0.01, 0.03] C: [-0.03, 0.01] D: [0.00, 0.03].

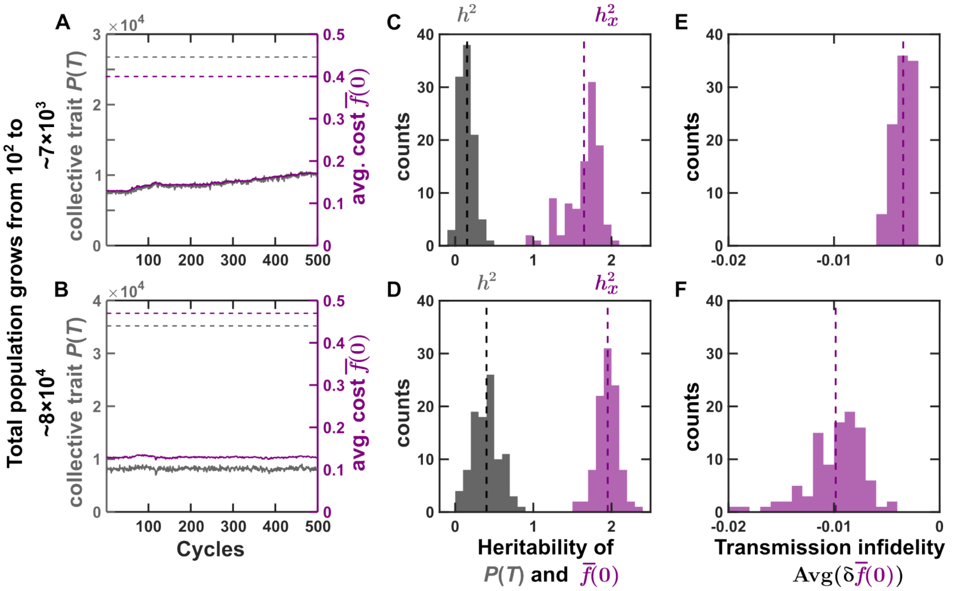


Figure S6: **Increasing the number of generations within each cycle of maturation improves heritability but not the selection outcome.** Results presented in the top row are from the simulation plotted in Figure 2D (Resource in each Newborn $R(0) = 1$, maturation time $T = 17$). Results presented in the bottom row are from simulation with $R(0) = 10$ and $T = 22$, which results in a higher number of generations and thus a larger microbial population in Adults within each cycle. (A, B) Evolutionary dynamics of chosen community trait $P(T)$ and heritable determinant $\overline{f}(0)$ over cycles. The gray and purple dashed lines correspond to the maximal $P(T)$ and the corresponding optimal $\overline{f}(0)$, respectively. A is identical to Figure 2D(i). (C, D) Both $h^2$ (heritability of the trait $P(T)$) and $h_x^2$ (heritability of the determinant $\overline{f}(0)$) are larger in D than in C. Histograms of the $h^2$ and $h_x^2$ are plotted in gray and purple, respectively. The means of the histograms are plotted in vertical dashed lines. The 95% confidence interval of each mean is calculated using the MATLAB bootci function with $10^4$ bootstrap samples. The mean $h^2$ and the corresponding 95% confidence interval in C and D are 0.15 ([0.14, 0.17]) and 0.40 ([0.37, 0.44]), respectively. The mean $h_x^2$ and the corresponding 95% confidence interval in C and D are 1.65 ([1.60, 1.69]) and 1.95 ([1.92, 1.98]), respectively. (E, F) The transmission infidelity (proportional to Avg $\left(\delta\overline{f}(0)\right)$, Eq. 12) is more negative in F than in E. This is because the fraction of non-producing M reaches a higher level with a higher number of generations during maturation. In total, the higher heritability gained by the strategy in the lower row is undercut by the more negative transmission infidelity, thus failing to improve the selection outcome.

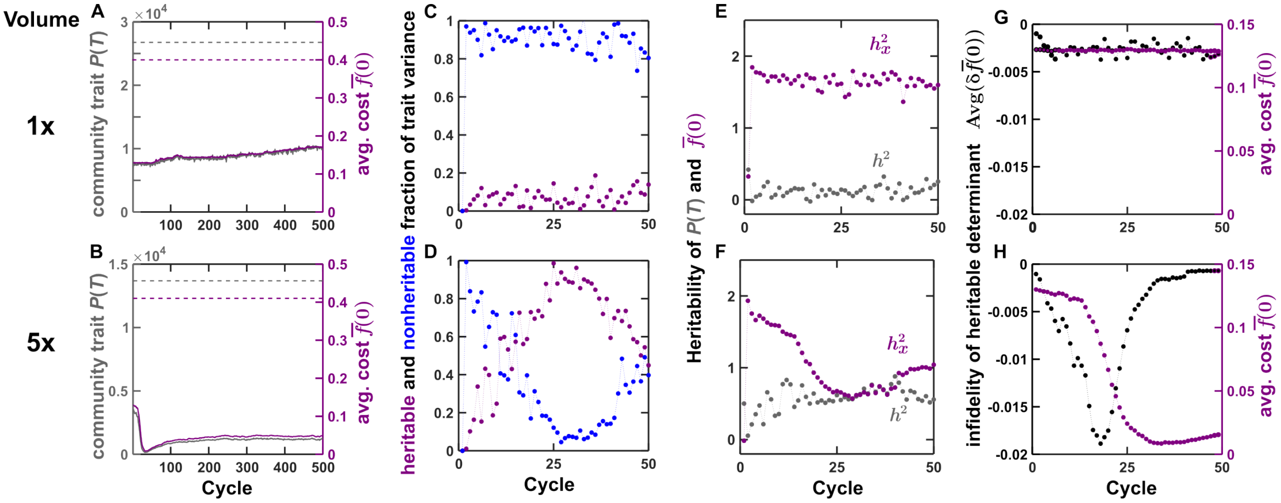


Figure S7: **Scaling up community volume and population size decreases nonheritable variation without improving the selection outcome.** Results presented in the top row are from the simulation plotted in Figure 2D (Resource in each Newborn $R(0) = 1$, the total population in each Newborn $BM(0) = 100$). Results presented in the bottom row are from simulation with $R(0) = 5$ and $BM(0) = 500$, equivalent to increasing the volume of communities by 5 times. (A, B) Evolutionary dynamics of chosen community trait $P(T)$ and heritable determinant $\overline{f}(0)$ over cycles. The gray and purple dashed lines correspond to the maximal $P(T)$ and the corresponding optimal $\overline{f}(0)$, respectively. A is identical to Figure 2D(i). (C, D) The larger $h^2$ in D can be explained by a larger fraction of heritable variation $(\text{Var}\left(\alpha\overline{f}(0)\right)/\text{Var}\left(P(T)\right))$, plotted in purple dots) and a smaller fraction of nonheritable variation $(\text{Var}\left(\alpha\phi_M(0)\right)/\text{Var}\left(P(T)\right))$, plotted in blue dots). (E, F) $h^2$ (heritability of the trait $P(T)$, plotted in gray dots) is larger in F than in E, but $h_x^2$ (heritability of the determinant $\overline{f}(0)$, plotted in purple dots) is smaller in F than in E. (G, H) Over the first 20 cycles, change in the heritable determinant $\overline{f}(0)$ due to intra-community evolution is much more negative in H than in G. This drives community trait $P(T)$ to nearly 0 as shown in B, despite higher heritability gained by the strategy in F.
